# Coordinated neural progenitor adaptations drive primate neocortical downscaling

**DOI:** 10.64898/2026.02.19.706795

**Authors:** Lidiia Tynianskaia, Cesar Mateo Bastidas-Betancourt, Esther Marie Grewe, Julia Marie Kniep, Neringa Liutikaite, Nesil Esiyok, Sabrina Heide, Nancy Rüger, Dimitri Lindenwald, Charis Drummer, Stoyan Petkov, Nataliya Di Donato, Michael Heide

## Abstract

Most primates possess a large, highly folded (gyrencephalic) neocortex, a feature present in the primate common ancestor. In contrast, several New World monkey species display a comparatively small and largely smooth, unfolded (lissencephalic) neocortex. One prominent example is the common marmoset, an increasingly popular model in neuroscience. This phenotype likely reflects an evolutionary reduction from the ancestral primate condition, implying modifications in neurodevelopmental programs. One essential basis for neocortical development is the activity and behavior of neural progenitor cells (NPCs). Here, we identify coordinated adaptations in NPC biology that bias neurodevelopmental trajectories toward neocortical downscaling. By combining marmoset and human cerebral organoids with analyses of fetal marmoset neocortical tissue and previously published histological data, we uncover multiple adaptations in apical and basal progenitors that converge on reduced progenitor capacity, alter early progenitor dynamics, and likely constrain neuronal output, thereby limiting the size and folding of the marmoset neocortex.

## Introduction

The neocortices of primates are generally large and highly folded (gyrencephalic), a feature commonly associated with advanced cognitive abilities^1–3^. Nevertheless, among New World monkeys (Platyrrhini), several primate species exhibit comparatively small and largely smooth (lissencephalic) neocortices, a feature not observed in Old World monkeys and apes (Catarrhini)^4^. One such species is the common marmoset (*Callithrix jacchus*), a small-bodied monkey widely used as a non-human primate model in biomedical research^5, 6^. Comparative and phylogenetic analyses suggest that the common ancestor of primates, and most likely also that of Platyrrhini, possessed a moderately sized, gyrencephalic neocortex^7^. Therefore, the largely smooth neocortex of the marmoset represents secondary lissencephaly^8, 9^, likely resulting from evolutionary dwarfing and the associated reduction in cortical size, despite the marmoset’s position within the otherwise predominantly gyrencephalic primate clade. As such, the marmoset can be viewed as an evolutionary exemplar of primate cortical size and folding reduction, providing a rare opportunity to dissect how a conserved primate neurodevelopmental program can be tuned toward reduced cortical expansion.

As a consequence, the marmoset exhibits the canonical neurodevelopmental cell type composition and cytoarchitecture characteristic of most primates ^10^, reflecting a developmental program that evolved in the context of a large and gyrencephalic neocortex. Cortical size and folding are closely linked to the behavior of neural progenitor cells (NPCs), the stem cells that generate neurons and glia. NPCs initially reside in the ventricular zone (VZ) as neuroepithelial cells (NECs), which undergo symmetric divisions to expand the progenitor pool before transitioning into apical radial glia (aRG)^11–13^. NECs and aRG are collectively called apical progenitors (APs). aRG can self-amplify through symmetric divisions or produce neurons and basal progenitors (BPs) via asymmetric divisions. In some species, such as most rodents, aRG generate neurons both directly and indirectly via BPs, whereas in primates, including marmoset, neuron production predominantly occurs indirectly through BPs^14, 15^. BPs migrate basally to populate the subventricular zone (SVZ) and are subdivided into basal intermediate progenitors (bIPs) and basal radial glia (bRG). bIPs are typically multipolar, undergo a limited number of divisions, and mainly produce neurons, whereas bRG retain a basal process, have higher proliferative potential, and contribute to both extensive further progenitor amplification and subsequent neuron production ^16–20^. The expansion and diversification of BPs, particularly bRG, leads to a pronounced expansion of the SVZ and its morphological subdivision into an inner and an outer SVZ (iSVZ and oSVZ), key features of neocortical development in species with large and highly folded neocortices^21, 22^. Strikingly, both a high abundance of bRG and an iSVZ/oSVZ separation are also observed in marmoset fetal brain, a clear contrast to the small and nearly lissencephalic appearance of its neocortex.

Accordingly, in the marmoset, adaptations must exist that adjust this neurodevelopmental program, that promotes high neuronal output in other primates, toward a lower neuronal output that results in a smaller and less folded brain. Several mechanisms have been proposed to explain this, including reduced amplification of BPs and bRG^14, 20^, lower diversity and proliferative potential of progenitors in the oSVZ^22, 23^, a bias toward neurogenic versus self-renewing divisions^17, 24^, and differences in progenitor cell-cycle dynamics, such as longer G1 or reduced proliferative capacity^25, 26^. Importantly, these mechanisms have not been experimentally analyzed in marmoset, and most evidence remains descriptive or inferred from comparative analyses, highlighting the need to investigate which progenitor features actually limit neuronal output.

Systematic *in vivo* analyses of marmoset NPCs are constrained by ethical considerations: the large number of animals required and limited (immediate) translational benefits make large-scale experimental studies difficult to justify. Brain organoids, however, offer a powerful alternative, modeling key aspects of NPC proliferation, differentiation, and spatial organization while reducing the need for extensive *in vivo* experimentation^27–30^. Shortly after the development of human brain organoids, protocols were established for brain organoids from non-human primates, mainly for great apes^31–33^. We recently published a unified protocol enabling the generation of cerebral organoids from human, chimpanzee, rhesus macaque, and marmoset induced pluripotent stem cells (iPSCs)^34^. This now allows the study of marmoset neocortex development using minimal tissue samples.

Here, we leveraged marmoset brain organoids alongside human organoids, marmoset neocortex tissue, and previously published histological data to investigate the cell biological features of marmoset progenitors that may limit neuronal output relative to most other primates. Using single-cell RNA sequencing (scRNA-seq), bulk RNA-seq of FACS-enriched progenitor populations, cell-cycle and immunofluorescence-based analyses of cell cycle, cellular composition, and progenitor properties, we identify multiple adaptations across both apical and basal progenitor types, regarding cell-cycle length, developmental timing, cleavage plane angle, and cell morphology. These findings provide mechanistic insight into how progenitor dynamics shape the small and largely unfolded marmoset neocortex, illuminating cellular strategies by which a conserved neurodevelopmental program can be evolutionarily tuned toward reduced neocortical size and folding.

## Results

### Marmoset and human cerebral organoids share cytoarchitecture and overall cell type composition but diverge in developmental timing and aRG features

To investigate the differences in cortical development that that underlie the formation of lissencephalic (marmoset) versus gyrencephalic (human) brains, we generated cerebral organoids from three independent human and three independent common marmoset induced pluripotent stem cell (iPSC) lines using our previously published protocol for primate cerebral organoid generation (Fig S1A)^34^. Despite their smaller overall size (Fig 1A, Fig S1B) and slightly faster neurodevelopmental pace during embryoid body stages (indicated by an enrichment of the GO term “nervous system development” in marmoset compared to human bulk RNA-seq at day 3 and day 6 (Fig S1C,D; Supplementary table 1)), marmoset cerebral organoids closely resemble human organoids under wide-field microscopy at 27 days of culture (Fig. 1A). This time point corresponds to the stage at which the typical cytoarchitecture of the early developing cortex is established in our organoid protocol^35, 36^. Immunofluorescence showed that cerebral organoids of both species predominantly acquire forebrain identity, as indicated by widespread FOXG1 expression (Fig. 1B). As expected for this developmental stage, SOX2-positive apical progenitors (APs) were localized near the ventricular surface, forming a ventricular zone (VZ), while the more basal regions were occupied by SOX2-positive basal progenitors and TUJ1-positive neurons (Fig. 1B). This organization reflects the typical apical-basal polarity of the developing cortical wall^37–39^, which is reflected in organoids of both species (Fig. 1B). These findings indicate that the fundamental cytoarchitecture of primate cerebral organoids is also conserved in marmoset cerebral organoids.

**Figure 1:**
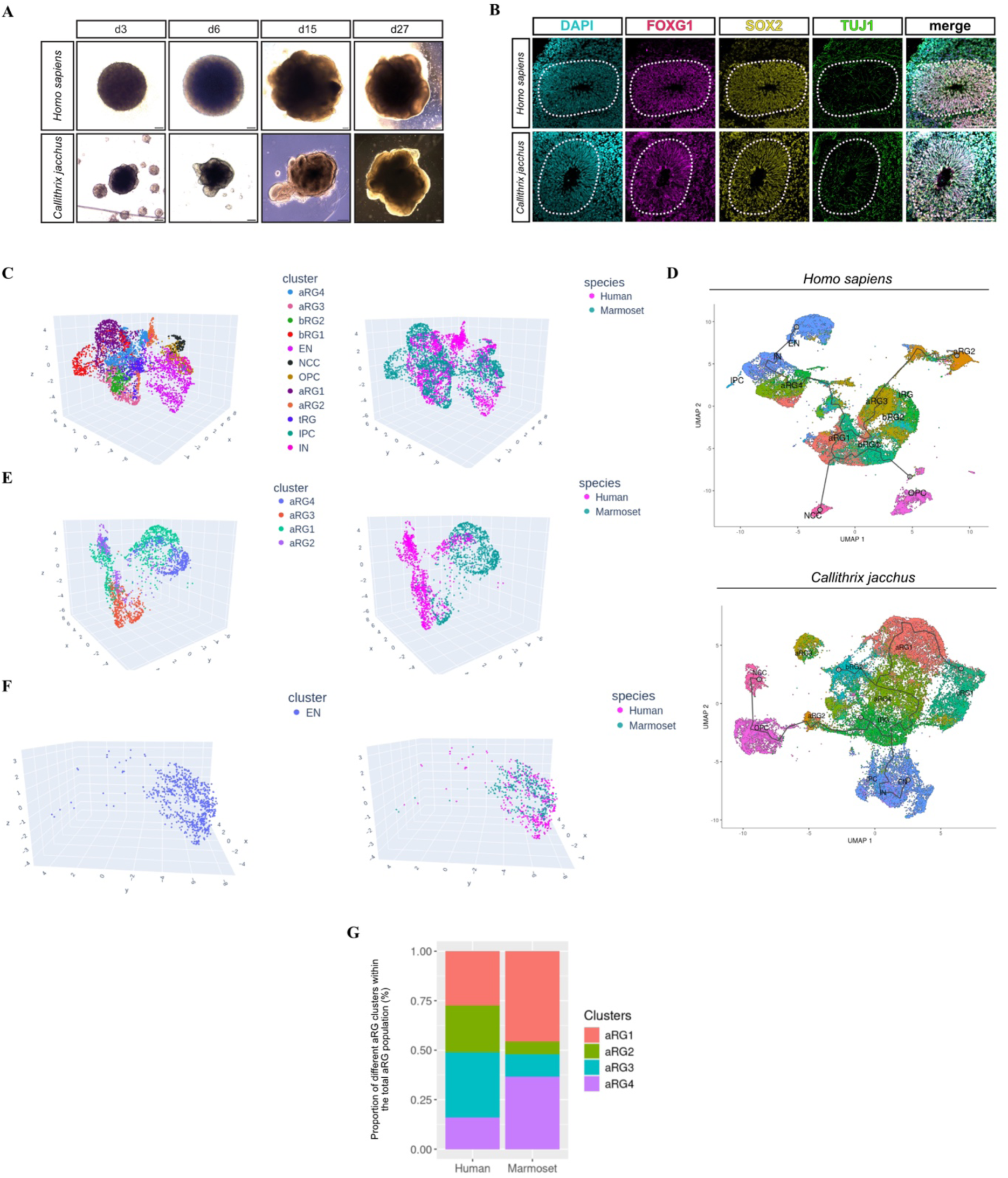
Comparable cytoarchitecture and cell-type composition but divergent apical radial glia cluster proportions in marmoset and human cerebral organoids. **(A)** Bright field images of human (upper row) and marmoset (lower row) embryoid bodies (3 and 6 days old) and cerebral organoids (15 and 27 days old). Scale bars, 100 µm (d3 and d6), 200 µm (d15 and d27). **(B)** Triple immunofluorescence for FOXG1 (magenta), SOX2 (yellow) and TUJ1 (green), combined with DAPI staining (cyan), of 30-day-old human (upper row) and marmoset (lower row) cerebral organoids. Dashed line indicates VZ/basal region border. Scale bars, 100 µm. **(C, E, F)** Three-dimensional uniform manifold approximation and projection (3D-UMAP) plot frames of single cells from 30-day-old human and marmoset cerebral organoids. Each dot represents one cell. Cells colored by transcriptionally defined cell clusters (left) and by species of origin (right; cyan, human; magenta, marmoset). Plots include 51,075 human cells and 46,667 marmoset cells from approximately 50 cerebral organoids in total. All identified cell types are shown in **(C)**, while **(E,F)** display subsets of the same UMAP embedding showing apical radial glia (aRG) **(E)** and excitatory neurons **(F)** only. **(D)** Monocle3 UMAP visualization of developmental trajectories in 30-day-old cerebral organoids derived from human (top) and marmoset (bottom) iPSCs, colored by cell type. **(G)** Relative contributions of distinct aRG clusters to the total aRG population in human and marmoset cerebral organoids at day 30. Abbreviations: aRG, apical radial glia; tRG, truncated radial glia; bRG, basal radial glia; IPC, intermediate progenitor cell; EN, excitatory neuron; IN, interneuron; OPC, oligodendrocyte progenitor cell; NCC, neural crest cell.

We next asked whether not only the basic cytoarchitecture but also the overall cell type composition is conserved in marmoset cerebral organoids compared to human. To address this, we performed single cell RNA sequencing (scRNA-seq) on day 30 cerebral organoids from both species and compared their cellular composition. This revealed the same cell populations in similar relative proportions in both species (Fig 1C, Fig S2A). Within this dataset, we identified four distinct aRG clusters, which could be distinguished by combinations of literature-defined marker gene expression (Supplementary table 2). Specifically, aRG1 expressed markers such as *HES5*, *HMGB2*, *TJP1*, *NUSAP1* indicative of a highly proliferative state; aRG2 expressed, among others, *SPARC*, *TCF12*, *PARD3B*, *TOP2A* likewise reflecting high proliferative activity; aRG3 expressed *TJP1*, *SPARC*, *TMEM14B*, *CLIC6*, *BMP7* and *COL9A2* pointing to a more differentiative state towards a basal progenitor fate; and aRG4 expressed *HES5*, *ZFP36L1*, *NPAS3*, *SLC1A3*, *ERBB4* pointing to a more differentiative state towards an astrocytic fate (Fig S2B).

To understand how these transcriptionally defined aRG states relate to lineage progression, we performed pseudotime trajectory analysis. This revealed clear species-specific differences in aRG dynamics. In human cerebral organoids, aRG clusters progressed along a relatively linear neurogenic trajectory, with multiple aRG clusters contributing to intermediate transcriptional states between aRG1 and neurons (Fig. 1D). In marmoset organoids, by contrast, the trajectory was less linear and more truncated, with fewer clusters contributing to intermediate states along the neurogenic pathway (Fig. 1D). Moreover, human aRG3 cluster extended over a longer pseudotime range, consistent with consistent with its expression of markers associated with neurogenic potential (Fig. S2C). In line with this, neurons reached the highest pseudotime values in human organoids. In marmoset organoids, however, glial cells—not neurons— occupied the highest pseudotime positions, suggesting earlier or more prominent glial specification (Fig. S2D,E). Together, these results indicate that marmoset cerebral organoids recapitulate the characteristically shorter neurogenic program of the marmoset relative to the human cortex, mirroring known *in vivo* differences (discussed in detail below) and highlighting that our organoid system captures species-specific developmental timing.

Having established that marmoset and human aRG differ in their lineage progression dynamics, we next examined whether these differences were also reflected in their transcriptional organization in UMAP space. When visualizing the scRNA-seq data by species, most clusters showed substantial intermingling of marmoset and human cells in the UMAP projection (Fig 1C). However, in the aRG clusters, a clear separation between marmoset and human cerebral organoid cells was observed (Fig 1D). This suggests that, despite overall similarity in cellular composition, marmoset and human aRG differ in their molecular profiles, which may reflect differences in aRG behavior or activity. In contrast, the excitatory neuron cluster did not show a separation by species (Fig. 1E), indicating that at these early stages of cerebral organoid development, species-specific differences are mainly confined to progenitor populations rather than more differentiated cells such as neurons. Moreover, when comparing the relative cell proportions of the aRG clusters only, we detected differences between marmoset and human (Fig 1F). Together, these findings indicate that while marmoset cerebral organoids, similar to the *in vivo* situation, recapitulate the overall cellular composition of human organoids at day 30, aRG show species-specific differences. This raises the possibility that already at the aRG level, developmental programs diverge between marmoset and human, potentially contributing to their distinct brain morphologies (small and lissencephalic versus large and gyrencephalic). Consequently, the next key question is which specific features of aRG differ between the two species.

### Slower cell cycle progression and reduced proliferative capacity of marmoset aRG underlie stable VZ/SVZ ratios in marmoset cerebral organoids and developing brain tissue

In early marmoset brain development, the relative proportions of the VZ to SVZ remains largely stable, with the VZ constituting the dominant germinal zone during much of neurogenesis. In humans, however, the SVZ rapidly expands and becomes the predominant germinal zone already at early stages of neurogenesis, reflecting early and robust production of basal progenitors, whereas in marmoset, basal progenitor production increases more gradually and peaks at later developmental stages^10^. Because basal progenitors are key contributors to cortical expansion and folding^40, 41^, this shift in timing of BP production may contribute to the formation of small, lissencephalic brains in marmosets.

This difference in timing of BP production raises the question of how VZ dominance is established and maintained in the marmoset. One possibility is that predominantly symmetric proliferative divisions, rather than asymmetric differentiative divisions, lead to reduced generation of basal radial glia (bRG) in favor of amplification of apical radial glia (aRG), consistent with early stages of cortical development. However, sustained proliferative divisions would also be expected to enlarge the VZ, which is not the case during marmoset development, suggesting that the proliferative capacity of aRG must be limited. One potential mechanism could be slower progression through the cell cycle. To test this hypothesis, we analyzed cell cycle length in marmoset and human cerebral organoids at culture day 30/31 using a cumulative EdU labelling approach^31, 42^ (Fig. 2A). We found that the total cell cycle length was longer in marmoset aRG (68.5 h) compared with human aRG (57.2 h) (Fig 2B,C) indicating that marmoset aRG indeed progress more slowly through the cell cycle and therefore generate fewer daughter cells.

**Figure 2:**
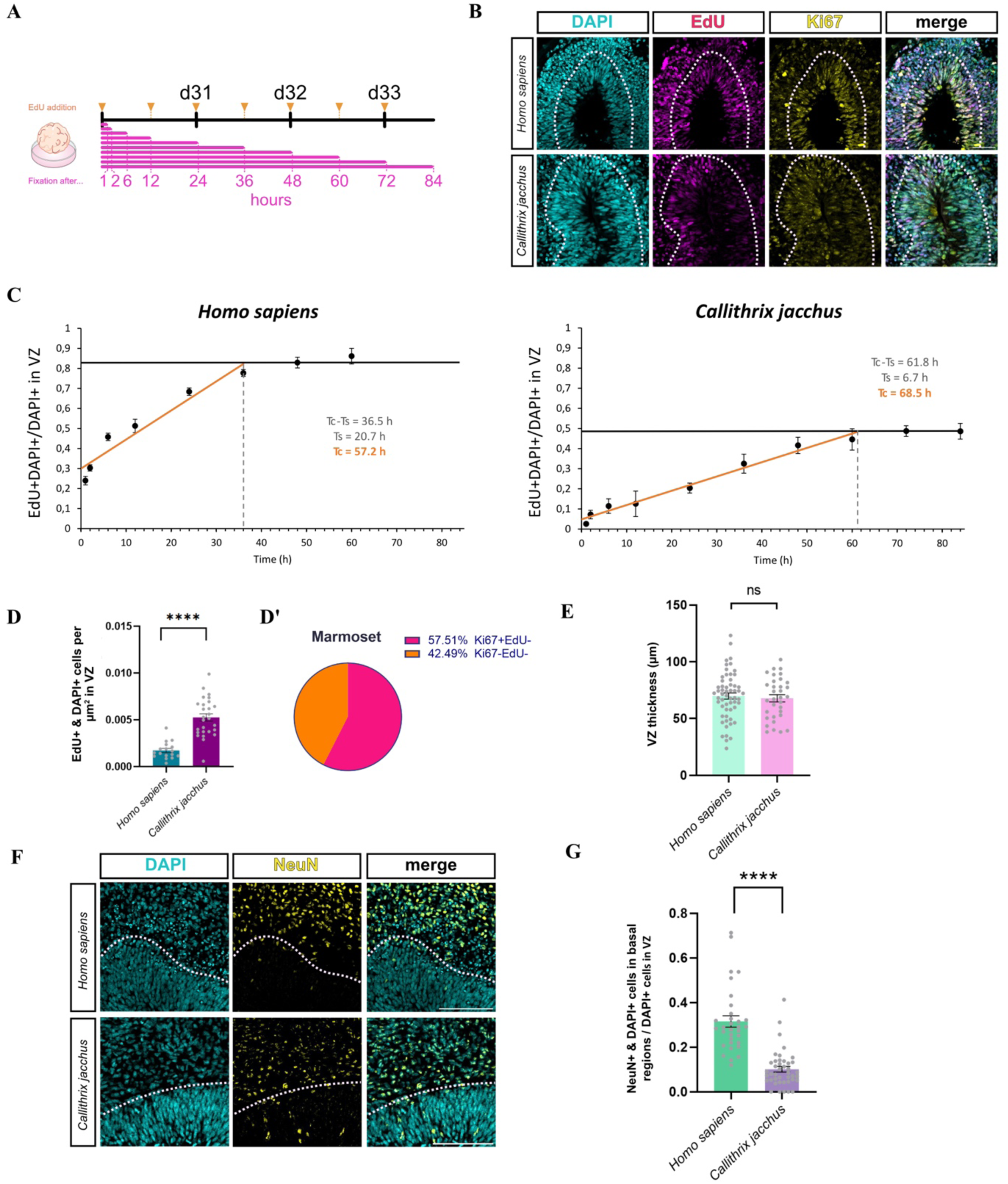
Slow cell cycle progression and low proliferative potential of marmoset APs allow VZ dominance to be maintained during much of neurogenesis. **(A)** Schematic timeline of the cumulative EdU labeling experiment. **(B)** Double immunofluorescence for EdU (magenta) and Ki67 (yellow), combined with DAPI staining (cyan), of a 32-day-old human (upper row) and 33-day-old marmoset (lower row) cerebral organoids subjected to EdU labeling at 30 days of culture for 48 h and 84 h respectively. Dashed line indicates VZ/basal region border. Scale bars, 50 µm. **(C)** Quantification of the proportion of DAPI-positive nuclei in the VZ which were also EdU-positive following 60 h (human) and 84 h (marmoset) cumulative EdU labeling in cerebral organoids. Linear regression lines for human (r² = 0,927) and marmoset (r² = 0,998) were used to estimate cell cycle parameters: Tc–Ts was read directly from the regression plots (vertical dashed lines). Ts was obtained from the x- and y-intercepts, and these values were then used to calculate the total cell-cycle length (Tc). Data represent the mean of multiple ventricle-like structures per time point from 1–2 independent batches, each comprising at least 3 cerebral organoids per iPSC line, generated from 2 independent iPSC lines per species (marmoset: cj_160419_5, cj_160419_6; human: SCTi003-A, CRTDi011-A). Error bars, ±SD. **(D)** Quantification of EdU+/DAPI+ double-positive cell numbers per µm^2^ in human and marmoset cerebral organoids under EdU-saturated conditions (48h and 60h of cumulative EdU labeling for human, 72h and 84h of cumulative EdU labeling for marmoset). Data represent the mean of 27 and 17 ventricle-like structures quantified in 10 common marmoset and 6 human cerebral organoids, respectively. Error bars, ±SEM; ****P < 0,0001 (Mann-Whitney test). **(D’)** Proportions of Ki67-positive and Ki67-negative cells among EdU-negative cells in marmoset cerebral organoids following 72h and 84h of cumulative EdU labeling analyzed in **(D)**. **(E)** Quantification of VZ thickness (µm) measured at four locations spaced 90° apart around a ventricle-like structure in 30-day-old human and marmoset cerebral organoids. Data represent the mean of 57 (human) and 33 (marmoset) ventricle-like structures derived from 16 human and 18 marmoset cerebral organoids, respectively, generated from 3 independent iPSC lines per species. Error bars, ±SEM; P=0,6553 (unpaired t-test). **(F)** Immunofluorescence for NeuN (yellow), combined with DAPI staining (cyan), in 30-day-old human (upper row) and marmoset (lower row) cerebral organoids. Dashed line indicates VZ/basal region border. Scale bars, 100 µm. **(G)** Quantification of NeuN+/DAPI+ double-positive cells in the basal region relative to DAPI+ VZ progenitors in the adjacent VZ region of 30-day-old human and marmoset cerebral organoids. Data represent the mean of 32 (human) and 40 (marmoset) radial units derived from 16 human and 16 marmoset 30-day-old cerebral organoids, respectively, generated from 3 independent iPSC lines per species. Error bars, ±SEM; ****P < 0,0001 (Mann-Whitney test).

Interestingly, during these cell cycle analyses we observed that the proportion of EdU-positive VZ cells plateaued at 83,9 % in human organoids but only at 48,7 % in marmoset organoids (Fig. 2C). This indicates that in marmoset cerebral organoids a large fraction of VZ cells did not incorporate EdU within the 84-hour labeling window, suggesting that they did not enter S-phase during this time. Thus, a substantial subset of marmoset aRG is either quiescent (i.e., exited the cell cycle) or proliferating at a very slow rate (i.e., reduced proliferative capacity). To investigate this further, we quantified the number of EdU-negative cells per VZ area in the samples past the point of saturation of the proliferative aRG population and found a significant increase in marmoset cerebral organoids compared to human (Fig 2B,D). Importantly, a bit more than half (57,5%) of these EdU-negative cells remained Ki67-positive (Fig. 2B,D’), indicating that this population consist of an actively (albeit slowly) proliferating population and a quiescent population.

We next asked whether the prolonged cell cycle and reduced proliferative capacity of marmoset aRG would affect VZ morphology and basal cell type (BPs and neurons) production in marmoset brain organoids. Quantification of VZ thickness in 30-day-old marmoset and human organoids revealed no significant differences (Fig. 2E), indicating that VZ thickness is established despite slower cell cycle progression of aRG. Moreover, quantification of NeuN-positive cells showed that in human cerebral organoids substantially more neurons are produced per VZ progenitor compared with marmoset organoids at day 30 (Fig. 2F,G) suggesting that reduced aRG proliferative capacity in marmosets contributes to the slowed down production of neurons.

Taken together, these data in marmoset cerebral organoids indicate that the VZ dominance observed in developing marmoset tissue is recapitulated in marmoset organoids and that it is based on the prolonged cell cycle and reduced proliferative capacity of aRG. This likely results in reduced production of BPs in early developing marmoset brain tissue^10^, which in turn contributes to the formation of the small, lissencephalic marmoset brain.

### Gene expression changes and cleavage plane orientation shifts underlie the late transition from VZ to SVZ dominance in marmoset brain development

In both marmoset brain tissue^10, 43^ and marmoset cerebral organoids (this study), the VZ is the dominant germinal zone during much of neurogenesis. However, as in other primates, marmoset neurons are produced mainly through indirect neurogenesis, i.e. via basal progenitors^14, 44^. Accordingly, the marmoset SVZ expands at later stages of cortical development into iSVZ and oSVZ, with a relative abundance of bRG comparable to that in ferret and human^10, 43^. Thus, at a certain developmental stage, the SVZ must replace the VZ as the dominant germinal zone. In marmoset brain tissue, this occurs as a sudden transition around GD85/86, when the SVZ rapidly expands^10^ and temporarily becomes the dominant germinal zone until neurogenesis ceases after GD10543. We therefore asked whether this abrupt shift from VZ to SVZ dominance is also recapitulated in marmoset cerebral organoids, and, if so, how it is regulated.

To address this, we first analyzed VZ thickness in marmoset cerebral organoids at culture days 30, 40, and 50 and found a significant reduction in VZ thickness (Fig. 3A,B). In parallel, the number of mitotic aRG decreased significantly (Fig. 3C) similar to the *in vivo* situation from GD78 to GD95^10^. By contrast, human cerebral organoids showed no reduction in VZ thickness in marmoset and maintained stable aRG mitotic activity between day 30 and day 50 (Fig. S3A–C). We next asked how this decline in VZ is mediated. A likely mechanism is increased delamination of aRG, resulting in more asymmetric differentiative and/or symmetric consumptive divisions. To test this, we dissociated marmoset cerebral organoids at culture days 30, 40, and 50, isolated aRG (high LeX, GLAST and PROM1 expression) by FACS using an approach similar to Johnson et al. 2015^45^ (Fig. S4A-B’), and compared their transcriptional profiles. To validate the enrichment of aRG, we examined TPM (Transcripts per Million) values of the FACS-isolated cells for established marker genes (see Methods) for aRG, bRG/BPs and neurons. The highest composite marker score was observed for aRG, confirming successful enrichment (Fig. S4D). Between day 30 and day 50, we detected decreased expression of NPC maintenance factors and of cilium assembly factors, accompanied by increased expression of ECM modulators and of neuronal differentiation factors (Fig. 3D; Supplementary table 3). Together, these transcriptional changes indicate a progressive increase in delamination signatures over time.

**Figure 3:**
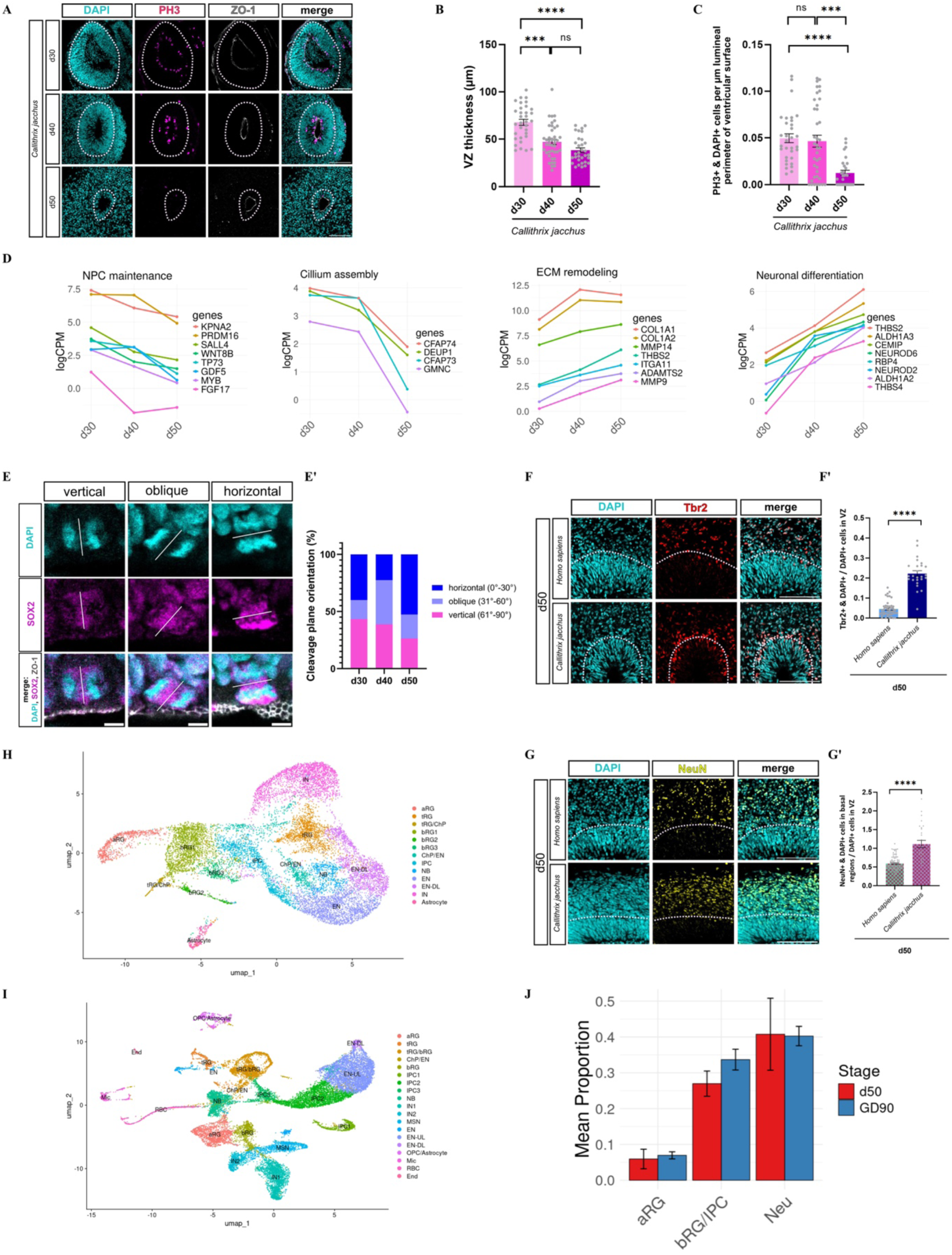
Coordinated changes in gene expression and cleavage plane orientation accompany AP delamination during marmoset cerebral organoid development and likely in vivo corticogenesis. **(A)** Double immunofluorescence for PH3 (magenta) and ZO-1 (gray), combined with DAPI staining (cyan), in 30- (upper row), 40- (middle row) and 50 (lower row)-day-old marmoset cerebral organoids. Dashed lines indicate VZ/basal region border. Scale bars, 100 µm. **(B)** Quantification of VZ thickness (µm) measured at four locations spaced 90° apart around a ventricle-like structure in 30, 40- and 50-day-old marmoset cerebral organoids. Data represent the mean of 33 ventricle-like structures from 18 marmoset cerebral organoids at day 30, 41 ventricle-like structures from 17 marmoset cerebral organoids at day 40, and 32 ventricle-like structures from 13 marmoset cerebral organoids at day 50, generated from 3 independent marmoset iPSC lines. Error bars, ±SEM; ns, not significant: P=0,1324; ***P=0,0001; ****P<0,0001 (Kruskal-Wallis test with Dunn’s multiple comparisons). **(C)** Quantification of PH3+ mitotic cells at the ventricular surface, normalized to luminal perimeter, in 30-, 40- and 50-day-old marmoset cerebral organoids. Data represents the mean of 32 ventricle-like structures from 18 marmoset cerebral organoids at day 30, 39 ventricle-like structures from 17 marmoset cerebral organoids at day 40, and 31 ventricle-like structures from 13 marmoset cerebral organoids at day 50, generated from 3 independent marmoset iPSC lines. Error bars, ±SEM; ns, not significant: P=0,634; ***P=0.0002; ****P<0.0001 (Kruskal-Wallis test with Dunn’s multiple comparisons). **(D)** LogCPM (log counts per million) plot of genes associated with NPC maintenance (left), cilium assembly (second from left), ECM remodeling (third from left) and neuronal differentiation (right), illustrating expression dynamics in aRG across marmoset cerebral organoid development at culture days 30, 40, and 50. **(E)** Double immunofluorescence for SOX2 (magenta) and ZO-1 (gray), combined with DAPI staining (cyan), in mitotic anaphase APs of 30- and 40-day-old cerebral marmoset organoids. Representative images show APs in anaphase with vertical (left; 30-day-old organoid), oblique (middle; 40-day-old organoid), and horizontal (right; 40-day-old organoid) cleavage plane orientations. White lines indicate the inferred axis of the cleavage plane, determined from the relative positions of the sister chromatids together with the local enrichment of SOX2 at the prospective cleavage site. Scale bars, 5 μm. The measured cleavage plane angles were 79° (vertical), 36° (oblique), and 10,9° (horizontal). **(E’)** Distribution of individual cleavage plane angles in anaphase APs from 30-, 40-, and 50-day-old marmoset cerebral organoids. Angles are grouped as vertical (61°–90°, magenta), oblique (31°–60°, light blue), and horizontal (0°–30°, dark blue). The dataset includes 30 (d30), 31 (d40) and 19 (d50) anaphase APs. **(F)** Immunofluorescence for Tbr2 (red), combined with DAPI staining (cyan), in 50-day-old human (upper row) and marmoset (lower row) cerebral organoids. Dashed line indicates VZ/basal region border. Scale bars, 100 µm. **(F’)** Quantification of the proportion of DAPI-positive nuclei in the VZ which were also Tbr2-positive cells in 50-day-old human and marmoset cerebral organoids. Data represent the mean of 45 ventricle-like structures from 11 human 50-day-old cerebral organoids and 25 ventricle-like structures from 11 marmoset 50-day-old cerebral organoids, generated from 3 independent iPSC lines per species. Error bars, ±SEM; ****P<0.0001 (Mann-Whitney test). **(H-I)** UMAP plots of scRNA-seq data from day 50 marmoset cerebral organoids (**H**) and GD90 marmoset telencephalic tissue (**I**), showing all annotated cell types. Plots include 14,438 cells from day 50 marmoset cerebral organoids and 23,052 cells from GD90 marmoset telencephalic tissue. aRG: apical radial glia, tRG: truncated radial glia, bRG: basal radial glia, IPC: intermediate progenitor cell, NB: neuroblast, EN(-DL or -UL): excitatory neuron (deep layer or upper layer), IN: interneuron, MSN: medium spiny neuron, ChP: choroid plexus, OPC: oligodendrocyte progenitor cell, NCC: neural crest cell, Mic: microglia, RBC: red blood cell, End: endothelium. **(G)** Immunofluorescence for NeuN (yellow), combined with DAPI staining (cyan), in 50-day-old human (upper row) and marmoset (lower row) cerebral organoids. Dashed lines indicate VZ/basal region border. Scale bars, 100 µm. **(G’)** Quantification of NeuN+/DAPI+ double-positive cells in the basal region relative to DAPI+ VZ progenitors in the adjacent VZ region of 50-day-old human and marmoset cerebral organoids. Data represent the mean of 99 (human) and 27 (marmoset) radial units derived from 19 human and 13 marmoset 50-day-old cerebral organoids, respectively, generated from 3 independent iPSC lines per species. Error bars, ±SEM; ****P<0.0001 (Mann-Whitney test). **(J)** Proportions of aRG, bRG/bIP, and neuron (EN+IN) clusters to the total cell population in marmoset day 50 cerebral organoids and GD90 telencephalic tissue. Data are the mean ± SD of 3 biological replicates per condition (∼30 cerebral organoids from 3 cell lines and 3 GD90 tissue samples).

Mechanistically, increased delamination by asymmetric differentiative and symmetric consumptive divisions has been linked to changes in cleavage plane orientation of aRG, from predominantly vertical to more oblique and horizontal. Such shifts increase the likelihood that one or both daughter cells will delaminate due to asymmetric inheritance of apicobasal cell components during division^46–48^. Consistent with this, analysis of cleavage plane orientation in aRG in anaphase revealed an increasing proportion of oblique and horizontal divisions from day 30 to day 50 in marmoset cerebral organoids (Fig. 3E,E’). Notably, the proportion of oblique and horizontal aRG divisions in marmoset cerebral organoids is already relatively high at day 30 (cumulative percentage: 56,5%), compared to published human data^31, 35, 49, 50^, mirroring in vivo observations in fetal marmoset brain tissue^43^. In this light, the prolonged cell cycle of marmoset aRG discussed in the previous paragraph likely further contributes to maintaining the VZ as the dominant germinal zone during early neurogenesis.

We next asked whether this change in cleavage plane orientation results in increased delamination and production of basal progenitors and neurons. To this end, we quantified TBR2-positive cells in the VZ, i.e. delaminated progenitors, and found an increased abundance in marmoset compared to human cerebral organoids (Fig. 3F,F’), indicating enhanced delamination. This was accompanied by an increase in TBR2-positive progenitors in basal regions, i.e. basal progenitors, between day 30 and day 50 in marmoset cerebral organoids (Fig. S5A,A’). Interestingly, quantification of HOPX-positive cells in basal regions revealed a small but significant increase in bRG, suggesting that the enhanced delamination in this time interval primarily gives rise to basal intermediate progenitors (bIPs) in marmoset cerebral organoids (Fig. S5B,B’). Finally, the proportion of NEUN-positive cells in basal regions, i.e. neurons, was higher in marmoset compared to human cerebral organoids (Fig. 3G,G’).

To corroborate these findings *in vivo*, we generated bulk transcriptome data from FACS-isolated aRG and neurons obtained from fetal marmoset telencephalic tissue at GD90 – a developmental stage at which the SVZ begins to emerge as the dominant germinal zone – using the same sorting strategy as for marmoset cerebral organoid samples but also including neurons/bIPs in G1 by using Vybrant DNA staining^51^ (Fig. S4A,C,C’). Marker-based analysis confirmed successful enrichment of aRG and, to a lesser degree, neurons from the tissue samples (Fig. S4E,F). We next asked which organoid stage most closely resembles the transcriptional state of the GD90 telencephalic tissue. To address this, we calculated the average Euclidean distance between the tissue-derived aRG and neuron transcriptomes and those of FACS-isolated aRG and neurons from marmoset cerebral organoids at days 30, 40, and 50. The shortest distances were consistently observed for day 50 organoids for both aRG and neurons (Fig. S4G,H), indicating that – from our available cerebral organoid samples – day 50 organoids most closely approximate the transcriptional state of GD90 marmoset telencephalic tissue. Following these results, we generated scRNA-seq datasets from day 50 marmoset cerebral organoids and from GD90 fetal marmoset telencephalic tissue (Fig. 3H, I and Fig. S6). To further validate that marmoset forebrain tissue more closely resembles marmoset cerebral organoids at day 50 than at day 30, we compared cell types across source (tissue vs. organoid) and stage and found that marmoset forebrain tissue at GD90 correlates better with marmoset cerebral organoid at day 50 than at day 30 (Fig. S7). By comparing day 30 and day 50 marmoset cerebral organoids, we detected a reduction in transcriptional complexity in day 30 organoids, with three distinct aRG clusters collapsing into a single cluster (Figs. 1C, 3H). In contrast, neuronal clusters displayed increased complexity, suggesting that as aRG diminish in both number and diversity, neuronal populations diversify. In addition, by comparing the relative proportions of aRG, bRG/bIPs and neurons (Neu) in day 50 cerebral organoids and GD90 telencephalic tissue, we found that, in line with the observed increase in delamination, in both datasets the mean proportions of BPs and neurons to total cells were higher than those of APs. Notably, the relative distributions of these three populations were highly similar between organoids and tissue (Fig. 3J). These scRNA-seq data are therefore consistent with a developmental stage in which increased aRG delamination contributes to a relative expansion of both BP and neuronal populations and further validates the shift to SVZ dominance in marmoset cerebral organoids at culture day 50.

In summary, these results demonstrate that the sudden expansion of the SVZ and concomitant decline of the VZ observed *in vivo* are recapitulated in marmoset cerebral organoids, albeit at different time points. Moreover, our data indicate that the sharp transition from VZ dominance to SVZ dominance in marmoset development is mediated by increased delamination, driven by changes in aRG gene expression and cleavage plane orientation. This late shift provides only a brief time window for BP amplification and neuron production before neurogenesis ends after GD105.

### Basal radial glia exhibit signatures of reduced proliferative capacity

After establishing the cell biological features of APs that likely underlie the small and lissencephalic marmoset brain, we next turned to basal progenitors (BPs). Similar to other primates, BPs in the common marmoset largely consist of basal radial glia (bRG), which are considered key contributors to the evolution of large, gyrencephalic brains due to their abundance and proliferative capacity^14, 15, 41^. Thus, mechanisms must exist in the marmoset to restrict neuronal output from bRG. One such mechanism is the short time window of SVZ dominance, which begins late and ends after GD105 in marmosets^10, 43^. This likely limits the time available for bRG proliferation and neuron production. However, given the intrinsically high proliferative capacity of bRG, this temporal restriction alone may not be sufficient, suggesting that additional, likely intrinsic mechanisms act to limit bRG proliferation in marmoset.

To identify such intrinsic mechanisms, we performed differential functional profiling of human and marmoset bRG using our day 30 human and marmoset cerebral organoid scRNA-seq dataset. We found that in comparison to marmoset, human bRG are enriched for modules related to cytoskeleton organization, extracellular matrix, and membrane biogenesis/assembly (Fig. 4A). These data are consistent with the concept that the proliferative capacity of bRG is associated with their morphology, specifically the number and orientation of their processes^16, 52, 53^. One important determinant of bRG morphology is PALMD. Previous work showed that increased PALMD expression correlates with more complex morphology, i.e. increased process number and diverse orientation, and increased proliferative capacity^52^. We analyzed PALMD expression in bRG of our human and marmoset day 30 cerebral organoid scRNA-seq dataset and found that marmoset bRG show lower PALMD expression than human bRG (Fig. 4B). In addition, analysis of relative PALMD expression in our scRNA-seq data from day 50 marmoset cerebral organoids and GD90 fetal marmoset telencephalic tissue showed that bRG express PALMD at significantly lower levels than neurons do, and at levels comparable to those of aRG (Fig 4C). This stands in strong contrast to previous findings in human, where PALMD is more highly expressed in bRG than in neurons or in aRG^52^, indicating that bRG morphology in marmoset may be less complex.

**Figure 4:**
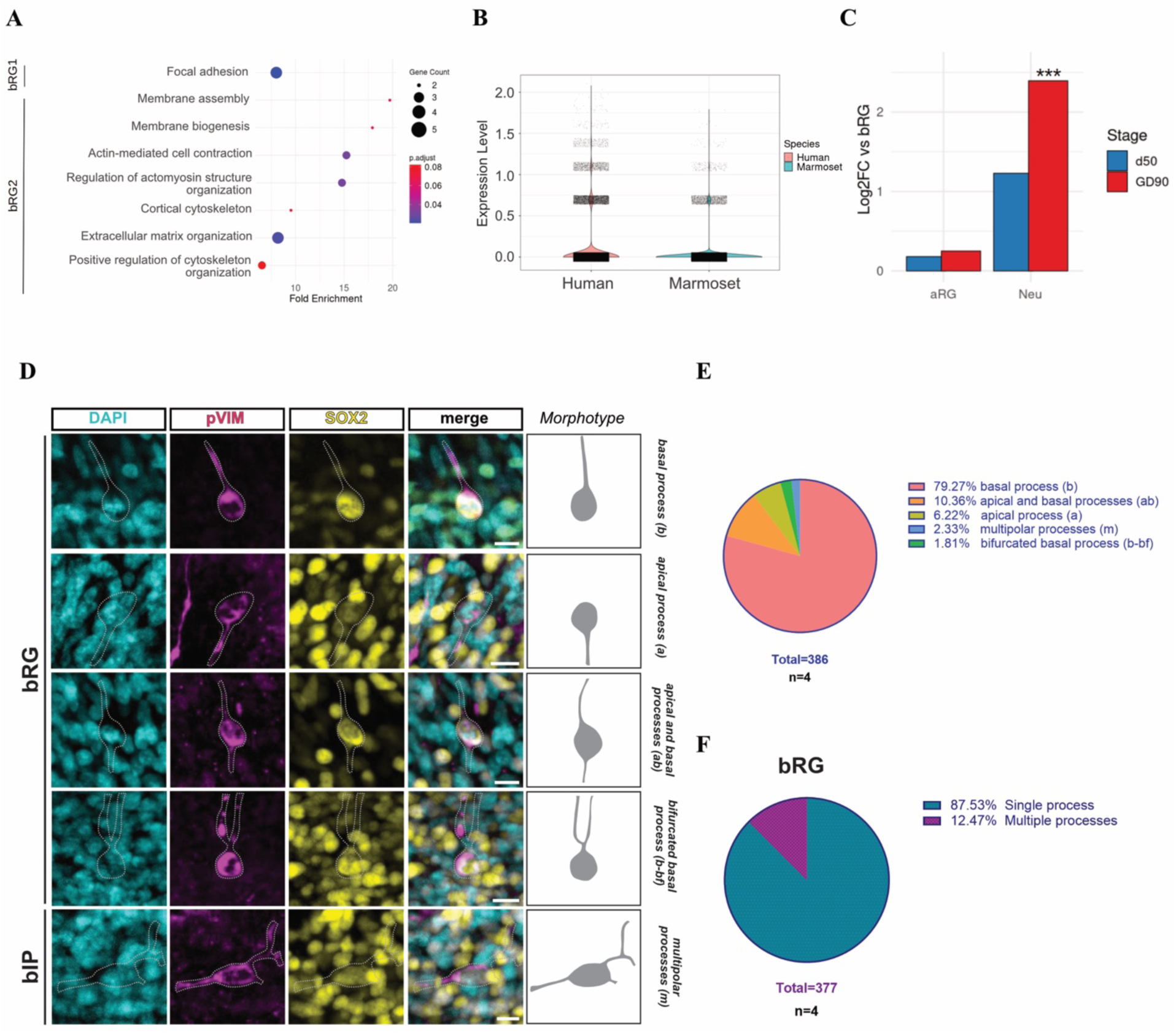
Marmoset BP gene expression profile and morphology indicate low proliferative potential in cerebral organoids and fetal telencephalic tissue. **(A)** Gene ontology enrichment analysis for Biological Process, Cellular Component and Molecular Function and KEGG (Kyoto Encyclopedia of Genes and Genomes) pathway analysis of genes enriched in bRG clusters of the scRNA-seq data of 30-day-old human and marmoset cerebral organoids. **(B)** Violin plot showing combined SCTransform-normalized expression of *PALMD* in bRG1 and bRG2 subpopulations of 30-day-old human and marmoset cerebral organoids. **(C)** Pseudobulk RNA-seq analysis comparing *PALMD* expression in aRG and neurons relative to bRG in day 50 marmoset cerebral organoid and GD90 telencephalic tissue. ***FDR<0.001. **(D)** Double immunofluorescence for pVIM (magenta) and SOX2 (yellow), combined with DAPI staining (cyan), in GD100 fetal marmoset cortex. Dashed lines indicate the outline of pVIM+ BPs. In all images, the apical side is positioned at the bottom. Representative images show pVIM+ BP morphotypes with basal process (b; upper row), apical process (a; second row from above), apical and basal processes (ab; third row from above), bifurcated basal process (b-bf; fourth row from above), and multipolar processes (m; lower row). Note that morphotypes b, a, ab and b-bf are associated with bRG, while m is associated with bIPs. Images were previously acquired for Heide et al., 2020^55^. Scale bars, 10 µm. (E) Distribution of individual pVIM+ BP morphotypes in GD100 fetal marmoset cortex. Morphotypes are grouped as described in (D) and based on the number of processes. Data represent the mean of 4 wild-type fetal cortices, with 16 frontal, 16 parietal, 16 occipital, and 8 temporal sections analyzed; images were previously acquired for Heide et al., 2020^55^ and re-analyzed here for morphological features.

To test this hypothesis, we next directly analyzed morphology BPs, including bRG. Because cerebral organoids do not fully recapitulate the abundance of bRG observed *in vivo*^27, 54^, we turned to in vivo tissue. In a previous study^55^, we performed phospho-vimentin (pVIM) immunofluorescence in wild-type and ARHGAP11B-expressing fetal marmoset neocortex at GD100. We re-analyzed the wild-type data, quantifying the number and orientation of processes of pVIM-positive cells in the SVZ. We classified mitotic BP into five morphological categories based on process number and orientation: single basal process (b, bRG), single apical process (a, bRG), one apical and one basal process (ab, bRG), single bifurcated basal process (b-bf, bRG) and more than two processes with various orientations (m, bIP) (Fig. 4C). This analysis revealed that the majority of mitotic BP extend only a single basal process, whereas mitotic BP with more complex morphologies are comparatively rare, especially when contrasted with previous data on human, rhesus macaque, and ferret (Fig. 4D,E). This indicates that bRG in marmoset neocortex exhibit less complex morphology in contrast to other primates.

In summary, in addition to the features of aRG and the shortened window of SVZ dominance, we identify adaptations in marmoset bRG toward reduced proliferative capacity. As bRG are thought to be major contributors to the expansion of large, folded brains, these adaptations likely contributed to the evolution of the small, lissencephalic marmoset brain from a gyrencephalic ancestor.

## Discussion

In this study, we used marmoset cerebral organoids together with human organoids, primary marmoset neocortex tissue, and re-analysis of published histological data to examine cell biological features of marmoset neural progenitors associated with a comparatively limited neuronal output. By integrating single-cell and bulk transcriptomic analyses with cell-cycle measurements and immunofluorescence-based analyses, we identify differences across progenitor populations, from apical to basal progenitors, that affect multiple aspects of progenitor behavior. These include changes in cell-cycle properties, developmental timing, cleavage plane orientation, and cellular morphology. Collectively, these features are consistent with VZ dominance during early marmoset brain development, an associated delay in basal progenitor production, and a reduced proliferative capacity of bRG that correlates with their simplified morphology. Rather than reflecting a single dominant mechanism, our results point to a combination of progenitor-level features that may collectively influence cortical growth in the marmoset. In the following sections, we discuss the usefulness of marmoset cerebral organoids in this study, how these distinct but interrelated parameters shape progenitor dynamics, their potential contribution to the small and largely unfolded marmoset neocortex, and the implications for secondary lissencephaly and primate cortical evolution.

To investigate the cell biological features of NPCs, we primarily relied on marmoset cerebral organoids. Throughout the analysis, however, we complemented these data with published *in vivo* studies of marmoset brain development, analyses of fresh fetal marmoset telencephalic tissue, and re-analysis of previously published histological datasets. This comparative approach enabled us to assess the extent to which marmoset cerebral organoids capture key aspects of marmoset cortical development. Notably, organoids recapitulate several cell biological features observed *in vivo*. One striking example is the temporal transition from VZ to SVZ dominance, which is reproduced in our cerebral organoids, albeit on a condensed timescale. We suggest that this correspondence may be facilitated by the an unguided/self-patterning organoid protocol that relies largely on intrinsic developmental programs rather than on extensive exogenous patterning cues, which could otherwise override endogenous timing mechanisms and obscure such temporal transitions. Together, these observations indicate that the marmoset cerebral organoid protocol used in this study provides a robust experimental system to investigate defined aspects of marmoset brain development and, for selected cell biological questions, may serve as a complementary or partial alternative to *in vivo* experiments in marmosets.

We focused on three main concepts that are likely to contribute to the small size and near-lissencephaly of the marmoset neocortex. First, we investigated the extent to which a relatively stable VZ-to-SVZ balance, characterized by a high VZ/SVZ ratio, is maintained during neurogenesis. Our analyses indicate that this VZ dominance is associated with an extended cell cycle length in APs, which maintains a low-turnover AP pool and reduces the frequency of BP production. Notably, a subset of VZ cells did not incorporate EdU within the analyzed labeling window, consistent with a very long cell cycle length. Among these cells, some were Ki67-positive, indicative of slow-cycling but proliferative progenitors, whereas others were Ki67-negative and may represent transiently quiescent cells, which has been described in rodents and was suggested for primates^56–60^. Differences in cell-cycle length between human and marmoset APs likely reflect species-specific modulation of rate-limiting steps in cell-cycle progression, such as G1/S commitment or S-phase progression. Interestingly, slower progenitor cell cycle dynamics and the delayed shift from VZ to SVZ dominance align with other aspects of marmoset development that proceed more slowly compared with other primates. These include prolonged Carnegie stages, later implantation relative to ovulation compared with rhesus macaques and humans, and a delayed transition from embryonic to fetal development^61–62^. Together, these observations suggest that progenitor dynamics may be embedded within broader changes in developmental timing, potentially reflecting secondary consequences of marmoset dwarfing, in which selective pressures on body size and life-history traits have led to non-uniform rescaling of developmental processes^63–64^.

Second, we investigated the late transition from VZ to SVZ dominance in marmoset brain development. This transition is recapitulated in marmoset cerebral organoids, albeit on a different timescale compared with *in vivo* development. We observed that it coincides with increased expression of extracellular matrix modulators and neuronal differentiation factors, together with decreased expression of neural progenitor maintenance factors and cilium assembly genes. These changes are consistent with increased progenitor delamination and expansion of the BP pool during the shift from VZ to SVZ dominance. Concomitantly, we detected a reduction in vertical cleavage plane orientation, which is predictive of a switch from symmetric proliferative to asymmetric differentiative divisions, further supporting BP expansion. Notably, both marmoset organoids (Fig. 3E’) and fetal marmoset neocortex^43^ display a lower proportion of vertical cleavage planes at early stages compared with other primates and humans^31, 35, 49, 50^, suggesting that marmoset APs may be biased toward asymmetric differentiative divisions from early stages onward. However, the combination of long cell cycle length and a subset of likely transiently quiescent APs may delay the full transition from VZ to SVZ dominance, preserving VZ dominance during early development. Third, we examined mechanisms that may limit the proliferative capacity of basal radial glia (bRG) in the marmoset neocortex. Despite its small and largely unfolded cortex, the developing marmoset neocortex contains a relative abundance of bRG comparable to that of other primates. This implies that mechanisms must act to constrain bRG expansion. One indirect constraint arises from prolonged VZ dominance and a delayed transition to SVZ dominance, which together reduce the temporal window available for bRG proliferation. Beyond this temporal restriction, we asked whether additional mechanisms acting more directly on bRG could further constrain their proliferative potential. One such feature is bRG morphology, as increased morphological complexity – reflected in the number, orientation, and branching of basal and apical processes – has been associated with higher proliferative capacity in gyrencephalic species, whereas simpler morphologies correlate with reduced proliferative behavior^16, 52, 53^. We find that marmoset bRG most frequently exhibit a simple morphology characterized by a single basal process (Fig. 4F). In contrast, more complex bRG morphologies, including multiple and branched processes, were reported at high frequency in humans and ferrets, representative of gyrencephalic mammals^52^. Together, these observations suggest that a bias toward simplified bRG morphology may represent an additional mechanism contributing to reduced bRG proliferative output in the marmoset, consistent with its limited cortical expansion and folding.

Together, our findings support a model in which multiple, partially overlapping adaptations across distinct progenitor populations contribute to the small size and near-lissencephaly of the marmoset neocortex. Although demonstrated here in the marmoset, these findings reveal a cellular strategy by which a conserved primate neurodevelopmental program may be evolutionarily tuned toward reduced cortical size and folding. Several of the features described here are expected to constrain bRG proliferative capacity, through reduced SVZ dominance, altered progenitor dynamics, and simplified bRG morphology; however, modulation of bRG alone appears unlikely to be sufficient. Support for this notion comes from an independent line of evidence: expression of the human-specific gene *ARHGAP11B*, which enhances bRG proliferative capacity via increased glutaminolysis^65^, in the fetal marmoset neocortex increases bRG abundance and neuronal output, resulting in an enlarged neocortex and induction of cortical folding, yet does not fully override the native developmental trajectory, as folding remains spatially restricted rather than affecting the entire neocortex^55^. This suggests that additional adaptations acting on apical progenitors, lineage progression, or developmental timing contribute at least equally to limiting cortical expansion in marmosets. In this view, neocortical size and folding are not simply lost in the marmoset lineage, but are actively constrained by multiple evolutionary modifications that counteract a primate-intrinsic developmental program favoring cortical expansion and folding. As a consequence, while the marmoset may be less well suited as a model to study the emergence of a large and highly gyrencephalic neocortex, it represents a particularly powerful system to investigate the cellular and developmental basis of secondary lissencephaly. In the present study, we focused on one central determinant of a small and nearly unfolded cortex reduced neuronal output but additional factors are also likely to contribute. These include mechanobiological influences, potentially shaped by differences in extracellular matrix composition or spatially patterned gene expression, such as genes differentially expressed in prospective gyri versus sulci or members of the FLRT family^66–69^. Future studies will be needed to determine whether such features can be modeled in cerebral organoids and whether they differ systematically between marmosets and primates with larger, gyrencephalic neocortices. Beyond these developmental mechanisms, recent work has suggested that glymphatic circulation is substantially more efficient in folded, gyrencephalic neocortex^70^. This raises the possibility that neocortical folding confers physiological benefits related to metabolic waste clearance, introducing an additional evolutionary pressure favoring gyrencephaly that may be relaxed in the context of the reduced brain size of the marmoset, consistent with its secondary lissencephaly.

## Methods

### Ethics

All animal experiments were conducted in accordance with the German Animal Welfare legislation (‘‘Tierschutzgesetz’’) and were approved by the local animal welfare authority, the Lower Saxony State Office for Consumer Protection and Food Safety (LAVES; 33.19-42502-04-23-00374). Common marmosets (Callithrix jacchus) were bred as pairs at the German Primate Center, Göttingen. Pregnancies were timed using established reproductive biology methods as previously published^71^. Each pregnancy was confirmed and subsequently monitored by ultrasonography. GD90 fetuses were retrieved by caesarean section performed by experienced veterinarians under sterile conditions and general anaesthesia, ensuring maternal survival, as previously described^71^. All animals received appropriate postoperative analgesic and antibiotic treatment.

### Cell lines

#### Induced pluripotent stem cell lines

All used iPSC lines were cultured under feeder-free conditions on Geltrex (0.16 mg/mL, Gibco, ThermoFisher)-coated dishes in the respective medium with daily medium changes. All human iPSC lines were kept at 37 °C and 5% CO_2_, while all marmoset iPSC lines at 5% O2, 5% CO_2_, and 90% N_2_ Human SCTi003-A (Stemcell Technologies) and CRTDi011-A^50^ were maintained in mTeSR1 (Stemcell Technologies), while iLonza2.2 iPSC cell line^72^ was cultured in StemMACS iPS-Brew XF (Miltenyi Biotec) supplemented with 1 μM IWR-1 (Sigma-Aldrich) and 0.5 μM CHIR99021 (Stemgent). All marmoset iPSC cell lines – cj_080619_4, cj_160419_5, cj_160419_6^73^ – were cultured in StemMACS iPS-Brew XF (Miltenyi Biotec) supplied with 3 μM IWR-1, 0.3 μM CGP77675 (Selleckchem), 0.3 μM Saracatinib (Selleckchem), 0.5 μM CHIR99021, 10 μM Forskolin (Selleckchem), 1 ng/mL Activin A (Sigma-Aldrich), 1 μM OAC1 (Selleckchem) as previously described^34, 73^. Versene (Gibco, Thermo Fisher Scientific) was used for routine splitting of all marmoset iPSC lines and iLonza2.2. Human SCTi003-A and CRTDi011-A were split using ReLeSR (Stemcell Technologies) according to manufacturer’s instructions. For iLonza2.2 and marmoset iPSC lines Pro-survival Compound (MerckMillipore) was used on the splitting day to boost cell survival.

### Generation of human and marmoset cerebral organoids

Human and marmoset iPSCs were differentiated into cerebral organoids following our established protocol^34, 35^. For EB generation on day 0, 9,000 cells per well of 96-well ultra-low attachment plates (Corning) were seeded in the respective iPSC medium supplemented with 50 µM Y-27632 (Abmole) or Pro-survival Compound. After 48 h, the medium was replaced with the respective one without Y-27632 or Pro-survival Compound. From day 4 for marmoset and day 4–5 for human, EBs were cultured in Neural Induction Medium consisting of DMEM/F12 (Thermo Fisher Scientific) supplemented with 1x N-2 supplement (Gibco, Thermo Fisher Scientific), 1x GlutaMAX (Gibco, Thermo Fisher Scientific), 1x MEM non-essential amino acids (Gibco, Thermo Fisher Scientific), and 1 µg/mL heparin (Sigma-Aldrich). The medium was exchanged every other day. On day 7 for marmoset and day 8–9 for human, EBs were embedded in Matrigel (Corning) and transferred to Differentiation Medium composed of a 1:1 mixture of DMEM/F12 and Neurobasal medium containing 0.5x B27 supplement without vitamin A (Gibco, Thermo Fisher Scientific), 0.5x N-2 supplement, 1x GlutaMAX, 0.5x MEM non-essential amino acids, 100 U/mL penicillin–streptomycin (PAN Biotech), 2.875 ng/mL insulin solution (Sigma-Aldrich), and 0.00035% 2-mercaptoethanol (Merck). Organoids were cultured on an orbital shaker at 55 rpm since then, with medium changes every other day. From day 13 for marmoset and day 14–15 for human onward, the medium was switched to Differentiation Medium containing B27 supplement with vitamin A (Gibco, Thermo Fisher Scientific) replacing its version without vitamin A and was changed every 3–4 days. From day 40 onwards, 0.1% Matrigel (Corning), 20 ng/ml brain-derived neurotrophic factor (BDNF; Peptotech) and 20 ng/ml neurotrophin 3 (Peprotech) were added to Differentiation Medium with B27 supplement with vitamin A. During all maintenance steps, cerebral organoids were kept in an incubator at 37°C and in a humidified atmosphere of 5% CO2 and 95% air.

### Cumulative EdU labeling

For labeling proliferating cells, EdU was added to 30–31-day-old cerebral organoids at a final concentration of 1 µg/mL from a 5 mg/mL EdU stock solution prepared in DPBS (Gibco, Thermo Fisher Scientific). The EdU-containing medium was exchanged every 12 h for a total labeling period of up to 60-84 h. Organoids were collected at defined time points (1, 2, 6, 12, 24, 36, 48, and 60 h for human organoids; with additional 72 and 84 h time points for marmoset organoids) and fixed in 4% paraformaldehyde (PFA; Merck) in DPBS (pH 7.5) for 15 min to 1 h at room temperature, washed three times with DPBS (Gibco, Thermo Fisher Scientific), and stored in DPBS at 4°C until further processing. EdU detection was performed using the Click-iT™ EdU Alexa Fluor™ 647 Imaging Kit (Invitrogen, C10340) according to the manufacturer’s instructions. Cell cycle parameters were calculated using linear regression analysis based on the model described by Nowakowski et al. 1989^42^.

### Immunofluorescence

Cerebral organoids were fixed in 4% PFA in DPBS (pH 7.5) for 15 min to 1 h at room temperature. Following fixation, organoids were washed three times in DPBS (Gibco, Thermo Fisher Scientific™) and stored in DPBS at 4°C until further processing. In preparation for cryosectioning, organoids were first sequentially incubated in 15% and 30% sucrose in DPBS overnight at 4°C, then embedded in cryomolds filled with TissueTek OCT (Sakura) and frozen on dry ice. Cryosections of 20 µm thickness were obtained using a CryoStar NX70 cryostat (Thermo Fisher Scientific), mounted onto SuperFrost Plus slides (Epredia), and stored at −20°C. Immunofluorescence staining was performed as previously described^35, 36^. In brief, sections were thawed and rinsed in PBS (pH 7.4). Antigen retrieval was performed in 0.1 M sodium citrate buffer (pH 6.0) for 1 h at 70°C, followed by cooling to room temperature. After washing in PBS, sections were permeabilized in 0.3% Triton-X100 in PBS (pH 7.4) and subsequently incubated in 0.1 M glycine in PBS (pH 7.4) for 30 min each. Blocking was carried out using 15% FCS in PBS for 30 min. Primary antibodies were diluted 1:200–1:300 in Can Get Signal® immunostain solution B (Toyobo) and applied overnight at 4°C. The following primary antibodies were used: FOXG1 (rabbit polyclonal, ab18259, Abcam), SOX2 (goat polyclonal, AF2018, R&D systems), TUJ1 (mouse monoclonal, 801201, Biolegend), Ki67 (rabbit polyclonal, ab15580, Abcam), NeuN (rabbit polyclonal, ab104225, Abcam), HOPX (rabbit polyclonal, PA5-90538, ThermoFisher Scientific), Phospho-histone H3 (rat monoclonal, ab10543, Abcam), ZO-1 (rabbit polyclonal, 61-7300, Thermo Fisher Scientific), Tbr2 (rabbit polyclonal, ab23345, Abcam). The slides were washed in 15% FCS in PBS and then incubated with secondary antibodies diluted 1:500 in Can Get Signal solution B (Toyobo) together with DAPI (1:1000) for 1 h at room temperature. The following secondary antibodies were used: Alexa Fluor 488: mouse (donkey polyclonal, A-21202, Invitrogen, Thermo Fisher Scientific), rabbit (donkey polyclonal, A-21206, Invitrogen, Thermo Fisher Scientific); Alexa Fluor 555: goat (donkey polyclonal, A-21432, Thermo Fisher Scientific), rabbit (donkey polyclonal, A-31572, Thermo Fisher Scientific), rat (goat polyclonal, A-21434, Invitrogen, Thermo Fisher Scientific); Alexa Fluor 647: rabbit (donkey polyclonal, A-31573, Invitrogen, Thermo Fisher Scientific). Slides were mounted with Mowiol (Carl Roth) and stored at 4°C in the dark.

### Image acquisition

Bright-field cerebral organoid images were acquired using a AX10 microscope (Zeiss) equipped with an Axiocam 105 color camera. Confocal images of immunostained sections were collected using a Zeiss LSM 800 microscope operated through the ZEN Blue software, equipped with 10×/0.45 M27, 20×/0.75 and 40×/0.95 Corr M27 air objectives. For larger fields of view captured as tile scans, individual tiles were automatically stitched using the Zeiss ZEN software. For Z stack acquisition, the images were usually taken at step intervals of 3.24 µm with the 10× objective, 2 µm with the 20× objective, and 1 µm with the 40× objective.

### Dissection and handling of marmoset fetal tissue

Isolated fetuses were decapitated, and the heads were immediately placed on ice. Fetal brains were dissected in ice-cold Hanks’ Balanced Salt Solution (HBSS), freed from the meninges, and the cerebral hemispheres were separated. From individual hemispheres, the telencephalon was further dissected. Dissected telencephalic tissue was subsequently dissociated for scRNA-seq and FACS.

### Dissociation of cerebral organoids and of fetal marmoset telencephalic tissue for scRNA-seq and FACS

For single-cell RNA-sequencing and FACS enrichment of aRG and neurons, tissue samples (150–300 mg, corresponding to ∼10 cerebral organoids or one GD90 fetal marmoset telencephalic tissue sample) were dissociated using a trypsin-based Neural Tissue Dissociation Kit (130-093-231, Miltenyi Biotec) according to the manufacturer’s protocol. Approximately 10 cerebral organoids were bisected and gently agitated in HBSS buffer to remove dead cells from the inner core. The organoids/tissue was then incubated with the enzymes indicated in the manufacturer’s protocol, stained with Trypan Blue, and counted using a hemocytometer, yielding ∼1 million cells collected in HBSS buffer.

### FACS enrichment of aRG and neurons from marmoset cerebral organoids and fetal telencephalic tissue

HBSS single-cell suspensions were resuspended in FACS buffer (PBS supplemented with 2% fetal bovine serum and 1 mM EDTA) and passed through a 100 μm strainer immediately before sorting. Approximately 300,000 cells were used for each isotype control and single-stain compensation gate, and the remaining cells were allocated for sorting. Staining was performed in FACS buffer at a 1:50 dilution for the GLAST-PE (mouse monoclonal, 130-118-344, Miltenyi Biotec) and LeX-FITC (mouse monoclonal, 130-113-484, Miltenyi Biotec) antibodies, and at a 1:10 dilution for the Prominin-APC (rat monoclonal, 130-123-793, Miltenyi Biotec) antibody. The corresponding isotype controls were IgG2a-PE (mouse monoclonal, 130-113-272, Miltenyi Biotec), IgMκ-FITC (mouse monoclonal, 553474, BD Pharmingen), and IgG1κ-APC (mouse monoclonal, 130-113-196, Miltenyi Biotec), each used at a 1:50 dilution. Cell suspensions were incubated with the antibodies for 30 minutes on ice, followed by three washes with FACS buffer. To assess DNA content in GD90 brain GLAST^low^/LeX^low^ cells, a Vybrant (Invitrogen, R37172) staining was performed at 37°C for 30 minutes after antibody washing.

FACS was performed on a Sony SH800 sorter with three lasers (488, 561, and 638 nm) using a 130 μm chip (LE-C3213, Sony), following the protocol described by Johnson et al. 2015^45^. Briefly, cells expressing GLAST and LeX (NPCs, GLAST^high^/LeX^high^) were further separated based on Prominin expression, which is high in aRG (Prom^high^). To isolate GD90 neurons, GLAST^low^/LeX^low^/Vybrant^low^ cells were collected (Fig. S4A,C,C’). Since neurons have exited the cell cycle, the peak with the lowest DNA content is enriched for neurons and bIPs in G1. Cells were sorted into 500 µL of lysis buffer (Micro RNeasy kit, Qiagen) prior to RNA extraction, as described below.

### RNA Extraction and cDNA Synthesis of FACS-enriched aRG and neurons and of EBs

RNA was extracted using the RNeasy Micro Kit (Qiagen) according to the manufacturer’s instructions, including on-column DNase I treatment. RNA was eluted in RNase-free water.

### Library preparation and sequencing

For bulk RNA-seq, two RNA-seq datasets were analyzed: (1) human and marmoset 3- and 6-day-old EBs, and (2) FACS-enriched cells from day 30, day 40 and day 50 marmoset cerebral organoids, as well as GD90 fetal marmoset telencephalic tissue (see above). RNA quality was assessed by measuring the RNA Integrity Number (RIN) using the Fragment Analyzer HS Total RNA Kit (DNF-472-FR; Agilent Technologies). Library preparation was performed on the STAR Hamilton NGS automation platform using the Illumina Stranded mRNA Prep Kit (Cat. No. 20040534) and the IDT for Illumina RNA UD Indexes Set A, Ligation with 96 Indexes (Cat. No. 20091646), starting from 200 ng of total RNA. The size distribution of the final cDNA libraries was determined with the SS NGS Fragment 1–6000 bp Kit on the Fragment Analyzer, yielding an average fragment size of 340 bp. Accurate library quantification was carried out using the DeNovix DS-Series System. Libraries were sequenced on a NovaSeq 6000 using an S2 flow cell (100 cycles), generating approximately 20–25 million reads per sample. For the analysis of 30-day-old human and marmoset cerebral organoids using scRNA-seq, cell suspensions were further processed using the 10x Genomics Chromium NextGem 3’ library preparation v3.1 kit, targeting 10–15k cells per sample. Libraries were sequenced on an Illumina NovaSeq 6000 S1 flow cell, aiming for 30,000 reads per cell (∼300 million reads per sample). Marmoset day 50 cerebral organoids and GD90 cortex scRNA-seq libraries were prepared using the Illumina PIP-seq Single Cell 3’ RNA Prep T10 Kit, following the manufacturer’s instructions (Cat. No. 2850744 and 20135691).

### Raw data QC and primary processing of bulk and scRNA-seq data

For bulk RNA-seq, raw sequencing signals were converted to BCL files using Illumina BaseCaller, followed by demultiplexing into FASTQ files with bcl2fastq v2.20.0.422. Read quality assessment and adapter trimming were performed using FastQC v0.12.1^74^.

For scRNA-seq, raw sequencing signals in BCL format were converted into FASTQ files using bcl2fastq2 (v2.20.0, 10x Genomics). Genome indexing and read alignment were performed using the CellRanger v8.0.0 count pipeline. Reads from human cerebral organoids were mapped to the GRCh38 human genome, whereas day 30 marmoset cerebral organoids were aligned to the human-to-marmoset lifted-off mCalJac1 genome annotation (Liftoff v1.6.3) provided by the ArchR package^75^. Day 50 marmoset cerebral organoid and GD90 cortical samples were aligned to the non-lifted-off version, as no cross-species comparison was intended. The CellRanger aggregate pipeline was then used to integrate the data for each species separately. Downstream analysis was performed using Seurat (v5.0.3)^76^, and high-quality cells were retained based on the following criteria: (1) gene expression counts greater than 500 UMIs, (2) number of detected genes per cell between 200 and 8,000, and (3) mitochondrial reads comprising less than 10% of total reads. A total of 97.742 30-day old human and marmoset cerebral organoid, 14.438 50-day old marmoset cerebral organoid, and 23.052 GD90 marmoset telencephalic cells passed these quality criteria and were included for further analysis. RNA-seq metadata is available in Supplementary table 4.

### Immunofluorescence- and image-based analyses

All quantifications were done blindly. Cell counts were performed in ZEN Lite or Fiji.

CEREBRAL ORGANOID DIAMETER QUANTIFICATIONS. Cerebral organoid diameter was quantified from bright-field images by averaging the length of two perpendicular lines drawn across one organoid.

CELL TYPE QUANTIFICATIONS. For cell type composition analysis, the number of DAPI-positive cells was quantified within VZ areas of ventricle-like structures. Subsequently, the number of cells positive for cell type–specific markers was determined in the adjacent basal area spanning 1.5–2 diameters of the quantified VZ zone corresponding roughly to the radial unit of the adjacent quantified VZ area. The number of cell type marker–positive cells was then normalized to the number of DAPI+ cells in the corresponding VZ zone, yielding one data point per radial unit in the presented graphs. This quantification approach effectively estimates cell type output per VZ progenitor. Mitotic cells in the VZ (mitotic APs) were quantified by measuring the luminal perimeter by ZO-1 staining and normalizing the number of PH3-positive cells adjacent to the ventricular surface to this perimeter. VZ radial thickness was measured from the ventricular surface (identified by ZO-1 staining) till the VZbasal-region boundary at four locations spaced 90° apart around a ventricle-like structure and averaged.

VZ–BASAL-REGION BOUNDARY AND VZ THICKNESS QUANTIFICATION. The boundary between the VZ and the basal region was identified based on differences in radial organization, nuclear density characteristic for VZ and in the initial analysis with the help of SOX2 immunofluorescence.

CLEAVAGE PLANE ORIENTATION. To assess cleavage plane orientation, mitotic anaphase APs were examined as previously described^35, 50^. The cleavage plane was inferred from the relative positions of the separating sister chromatids by examining z-stacks of DAPI-stained optical sections spanning the prospective division plane. The ventricular surface was identified by ZO-1 immunostaining. Cleavage plane angles were measured relative to the ventricular surface. Anaphase cleavage plane angles were categorized into three groups: vertical (90–61°), oblique (60–31°), and horizontal (30–0°).

STATISTICAL ANALYSES. All statistical analyses were performed in Prism (GraphPad Software). Outliers were identified using the ROUT method (Q = 1%). Normality of the data was evaluated using the Anderson– Darling, D’Agostino–Pearson, Shapiro–Wilk, and Kolmogorov–Smirnov tests. For comparisons between two groups, either an unpaired t-test or a Mann–Whitney test was applied depending on distribution. For analyses involving more than two non-normally distributed groups, a Kruskal–Wallis test followed by Dunn’s multiple-comparison test was used. All used statistical tests were two-sided.

### Transcriptome-based analyses

#### BULK RNA-SEQ ANALYSIS

All reads were mapped using STAR (v2.7.11b)^77^. Human reads were aligned to the GRCh38 genome, and marmoset reads were aligned either to the previously described human-to-marmoset lifted-off mCalJac1 genome annotation (EB data) or to the non-liftoff annotation (FACS-enriched cells). Mapping was performed using STAR with the alignment settings --alignEndsType Local, -- outFilterMatchNmin 20, and --outFilterMatchNminOverLread 0.3. Gene-level counts were generated using STAR’s --quantMode GeneCounts option. Differential expression analysis for both datasets was conducted using the edgeR package (v3.42.4)^78^. For the EB dataset, genes were considered differentially expressed if they showed an absolute log2 fold change > 1 and a false discovery rate (FDR) < 0.01. Gene ontology functional enrichment analysis of differentially expressed genes was performed using ShinyGO (v0.76.3)^79^ .

TPM values for well-established markers of aRG (PARD3, SOX2, HES1, HES5, PROM1, CENPF, MKI67, TJP1, TJP2, CDH2, SLC1A3, CRYAB, PALS, NUSAP, ASPM, KNL1), bRG/bIP (EOMES, NEUROG2, INSM1, BCL11A, PPP1R17, PTPRZ1, LIFR, FEZF2, PDGFD, BMP7, HOPX, TMEM14B), and neuronal populations (DCX, TUBB3, MAP2, RBFOX3, NEUN, SYN1, STMN2, BCL11B, SATB2, SLC17A7, SLC17A6) were quantified across all FACS-enriched populations to assess sorting purity. To determine which cerebral organoid stage most closely recapitulates GD90 fetal marmoset telencephalic cell populations, Euclidean distances were computed using the expression of 728 neurodevelopmental genes (Supplementary table 5) between aRG of day 30, day 40 and day 50 cerebral organoid and FACS-enriched GD90 fetal marmoset telencephalic aRG and neuronal samples.

#### SCRNA-SEQ DATA PROCESSING AND CELL ANNOTATION

To generate a single object comprising all data, Seurat’s SCTransform (v0.4.1)^80^ normalization pipeline was used to integrate the human and marmoset day 30 cerebral organoid data, first within each species and then across the two species. Harmony (v1.2.4) integration was applied to the SCT assay, retaining the first 19 PCA components for further analysis based on elbow plot assessment. Clusters were identified at a resolution of 0.27, determined using Clustree cluster analysis, resulting in a total of 17 clusters, which were subsequently marker-annotated using the standard Seurat downstream pipeline. Cell annotation was performed using a custom semi-automatic approach based on top markers described by Kanton et al. (2019)^32^, Braun et al. (2023)^81^, Polioudakis et al. (2019)^82^, and Hendriks et al. (2024)^83^ (Supplementary Table 2). Briefly, the top 10, 30, and 50 markers were intersected separately with reference markers to assign each cluster to a possible cell type. Cell cycle information was independently provided by Seurat’s CellCycleScoring function for each cluster and was used to distinguish potentially overlapping cell types (e.g., early versus late-stage aRG). Some clusters were subsequently merged into the same cell type, resulting in a final total of 13 cell types (Dotplot – Fig. S2B). Cells that were not annotated to any cell type were excluded from the cell proportion comparison (cluster 15: 387 human and 169 marmoset cells). Trajectories and pseudotime progression were inferred using the standard Monocle3 pipeline (v1.4.26)^84^. While pseudotime values per cluster per species were obtained by combining trajectory inference for human and marmoset cells, trajectory topography reconstruction was performed separately for each species. Briefly, dimensionality reduction was performed using PCA on the first 19 PCs, and cells were aligned across species using the align_cds function. Reduced-dimensional embeddings were computed with UMAP, and clustering was performed at a very low resolution (1e-6) to preserve fine-scale structure, followed by graph learning and cell ordering, rooted at aRG1. For the marmoset 50-day old cerebral organoids and GD90 fetal marmoset telencephalic tissue, the same integration, clustering, and cell type annotation pipelines were applied, selecting 26 and 25 PCs, respectively, and a resolution of 0.45 for both datasets. A total of 13 and 22 clusters were identified, corresponding to 13 and 19 cell types, respectively. Markers for annotated cell types are available in Supplementary table 6 for all scRNA-seq datasets.

#### PSEUDOCELL-BASED CROSS-DATASET CORRELATION ANALYSIS OF MARMOSET SCRNA-SEQ DATA

To enable robust comparisons across samples while reducing technical noise, dropout, and computational burden, pseudocells were generated for all marmoset single-cell RNA-seq data (day 30 and day 50 marmoset cerebral organoids, and GD90 fetal marmoset telencephalic tissue) by grouping individual cells based on their similarity in PCA space. First, the raw count matrix was extracted from the integrated Seurat object and merged into a sparse genes × cells count matrix. Principal component analysis (PCA) was performed on the integrated data, using the same number of PCs as in the original analysis to define the embedding.

A set of seed cells was randomly selected from the full cell population to serve as centers for grouping. Each cell was then assigned to the nearest seed cell based on Euclidean distance in PCA space, forming non-overlapping groups of cells. A sparse cell–pseudocell membership matrix was generated, and counts were aggregated and normalized by pseudocell size. The dominant biological identity (cluster and sample origin) was preserved for each pseudocell. The resulting pseudocell count matrices were further reduced to the common genes included across the three marmoset single-cell RNA-seq datasets (13,846 genes). Cell type–specific gene expression profiles were generated by aggregating pseudocell count matrices. Matching cell IDs between expression matrices and metadata were retained, and mean expression per gene was computed within each cluster. Pairwise Pearson correlations were then computed and plotted between all cell types across stages.

#### CELL TYPE-SPECIFIC DIFFERENTIAL EXPRESSION AND NETWORK ANALYSIS OF BRG

To investigate functional differences between human and marmoset bRG, we employed two complementary strategies: pseudobulk differential expression analysis and cell type–specific network modeling. First, single-cell transcriptomes from bRG clusters (bRG1 and bRG2) were aggregated into pseudobulk profiles using Seurat’s default AggregateExpression pipeline. Genes with a log2 fold change > 0.5 and an FDR-adjusted p-value < 0.05 were considered significantly differentially expressed. In parallel, we reconstructed and compared cell-type-specific gene networks across species using a multi-step approach. Single-cell RNA-seq data were first subsetted and harmonized by cluster in a Seurat object. The ACTIONet (v2.0.18)^85^ package was then applied to perform dimensionality reduction and latent network inference, generating cluster-level feature specificity scores. These scores were used by the SCINET framework (v1.0)^86^ to infer weighted cell-type-specific gene networks for each species. Human and marmoset networks were subsequently aligned and integrated using the scHumanNet (v0.1.0)^87^ v3 reference interactome, which provides a curated map of known human gene interactions, enabling meaningful cross-species comparisons. Node centrality was computed for each network, and composite hub scores were calculated across species. The top 50 hub genes per cluster were extracted, and species-specific hub genes were identified using a non-parametric approach (FindDiffHub, BH-adjusted p < 0.05), requiring a minimum of 15 cells per group. The intersection of the genes between both approaches that were found to be upregulated/more central for each cluster in each species was interrogated in ClusterProfiler v4.16.0^88^ functional profiling of GO:BP, GO:CC, GO:MF and KEGG modules. This integrative approach enabled us to pinpoint both transcriptional and topological divergences in progenitor cell identity across primate species.

#### *PALMD* EXPRESSION ANALYSIS

*PALMD* normalized expression values for bRG (bRG1 and bRG2) were directly compared between human and marmoset day 30 cerebral organoid cells from the integrated Seurat object, since *PALMD* was a highly variable gene (HVG) and therefore retained during downstream analysis. For 50-day-old organoids and GD90 cortex scRNA-seq, a pseudobulk analysis was performed to identify differentially expressed genes (DEGs) between cell types, with the contrasts being EN–bRG and aRG–bRG. Cell-level count data from the Seurat object were extracted, combined, and aggregated by cluster group, resulting in a pseudobulk expression matrix (genes × pseudobulk clusters). Differential expression analysis was conducted using the edgeR package, with the aforementioned biological replicates included as covariates.

## Data availability

Raw scRNA-seq and bulk RNA-seq sequencing data generated in this study have been deposited in the ENA under accession number PRJEB107058. Processed data, including gene expression matrices, Seurat objects and metadata used for downstream analyses, are publicly available in Zenodo (DOI: 10.5281/zenodo.18480599): https://zenodo.org/records/18480599?token=eyJhbGciOiJIUzUxMiJ9.eyJpZCI6ImJmYWViM2MzLTZmNDUtNDY0ZS04ZmRjLTE2ZTBmYzdmZDIxMSIsImRhdGEiOnt9LCJyYW5kb20iOiI4YWNlYmMxMGM1ZTcwNWU0NGE5ZGY3ZjY2YjdiOGRhOSJ9.z24aRNPMYvVLQGF9OIyTli5sJycxRxUZjzb33-ODqziW3U-TC4DYs7GlZleSM7HWegti5En3GQ0XOC4p99iE9g

## Code availability

All custom scripts and computational workflows used for data preprocessing, analysis, and figure generation are openly available at: https://github.com/MateoBastidasBetancourt/Tynianskaia_et_al_2026

## Acknowledgements

We apologize to all researchers whose work could not be cited due to space limitations. We thank G. Salinas and her team at the NGS Integrative Genomics (NIG) facility at the University Medical Center Göttingen (UMG) for bulk and single-cell RNA sequencing services and bioinformatic support. We thank A. Dahl and his team at the DRESDEN-concept Genome Center for single-cell RNA sequencing services and bioinformatic support. We also thank W. B. Huttner for granting access to previously acquired images of GD100 fetal marmoset neocortex sections published in Heide et al. (2020), and E. Sasaki and H. Okano for providing the original GD100 fetal marmoset neocortex tissue from which these images were obtained. We further thank R. Behr for access to personnel and infrastructure and for providing marmoset iPSC lines. In addition, we thank the members of the Heide lab for helpful discussions and for critically reading the manuscript. This work was supported by a DFG grant (DI 2170/5-1) to NDD, an ERC Starting Grant (PRIMAZINC, 101039421) to MH, and generous support from the Daimler and Benz Foundation (32-04/24) to MH.

## Author contributions

LT: conceptualization, methodology, investigation, formal analysis, visualization, writing – original draft, review, editing

CMB: conceptualization, software, investigation, formal analysis, visualization, data curation, writing – original draft, review, editin

EMG: Investigation, formal analysis

JMK: Investigation, formal analysis

NL: Investigation NE: resources

SH: investigation

NR: resources

DL: resources

CD: resources

SP: resources

NDD: funding acquisition

MH: conceptualization, supervision, project administration, funding acquisition, writing – original draft, review, editing

## Materials & Correspondence

Further information and requests for resources and reagents should be directed to the corresponding author, Michael Heide.

