## Supplementary Information for "Coordinated neural progenitor adaptations drive primate neocortical downscaling"

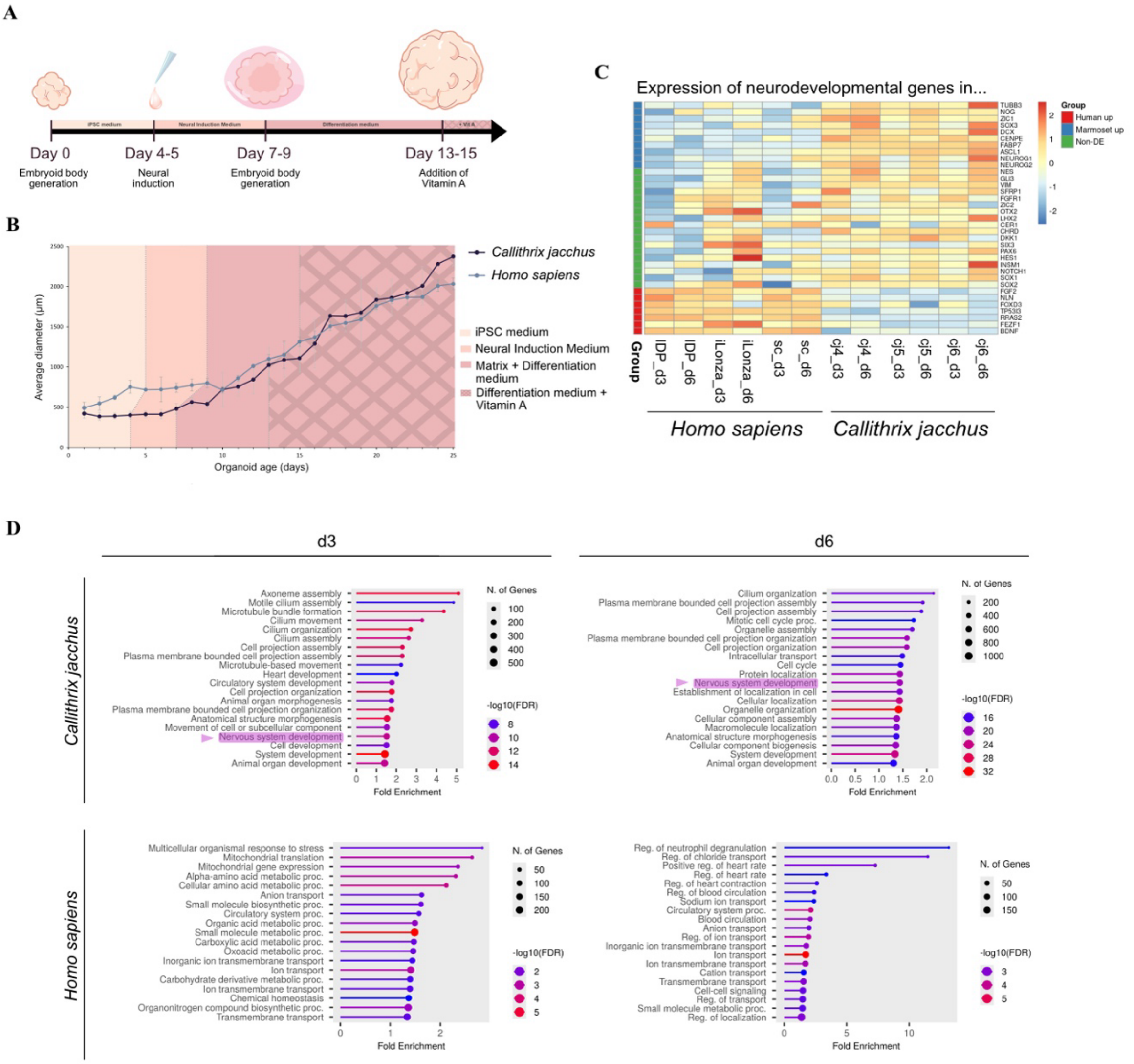

**Fig. S1: Marmoset embryoid bodies display smaller overall size but accelerated neural fate acquisition compared to human.**

**(A)** Schematic illustration of the timeline of cerebral organoid generation. **(B)** Growth curve showing the diameter of human and marmoset cerebral organoids analyzed over a 25-day period. Data represent the mean of 1–3 independent batches per time point, each comprising from 7 to 21 cerebral organoids (7 organoids per iPSC line) generated from 3 independent iPSC lines per species (marmoset or human). Error bars, ±SD. **(C)** Heatmap depicting differential gene expression of neurodevelopmental genes between human (left) and marmoset (right) embryoid bodies (EBs) at day 3 and day 6, based on three biological replicates (cell lines) per condition (species × stage). Genes are grouped by expression pattern: upregulated in human (red), upregulated in marmoset (blue), or non-differentially expressed (green). Differentially expressed genes were defined as those with $|\log_2 FC| > 1$ and $p < 0.01$. **(D)** Gene Ontology (GO): Biological Process enrichment analysis of genes overexpressed in marmoset (top) and human (bottom) EBs at day 3 (left) and day 6 (right).

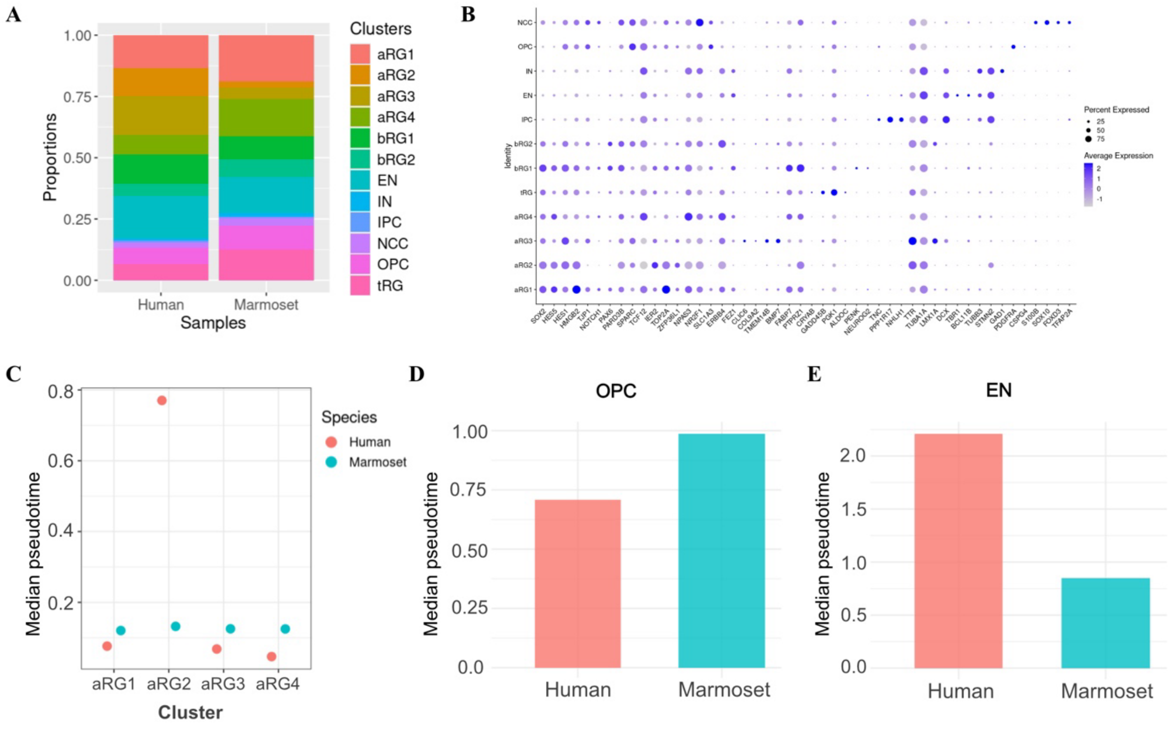


**Fig. S2: Marmoset and human 30-day-old cerebral organoids share cell type composition but differ in developmental maturation of aRG and terminal cell types.**

**(A)** Relative contributions of the various cell type clusters to the total cell count in human and marmoset cerebral organoids at day 30. aRG: apical radial glia, tRG: truncated radial glia, bRG: basal radial glia, IPC: intermediate progenitor cell, EN: excitatory neuron, IN: interneuron, OPC: oligodendrocyte progenitor cell, NCC: neural crest cell. **(B)** Dot plot of SCTransform-normalized scRNA-seq expression data showing average expression levels (color) and the percentage of cells expressing selected marker genes as indicated (dot size) in 30-day-old human and marmoset cerebral organoids. **(C–E)** Median pseudotime of 30-day-old human and marmoset cerebral organoids calculated using the Monocle3 standard pipeline, with all cell types rooted to aRG1 (see Methods). Pseudotime distributions were non-normal (Shapiro–Wilk test, $P < 0.05$). Only crossspecies comparisons of median pseudotime for aRGs **(C)**, oligodendrocyte precursors (OPCs; **D**), and excitatory neurons (ENs; **E**) are shown.

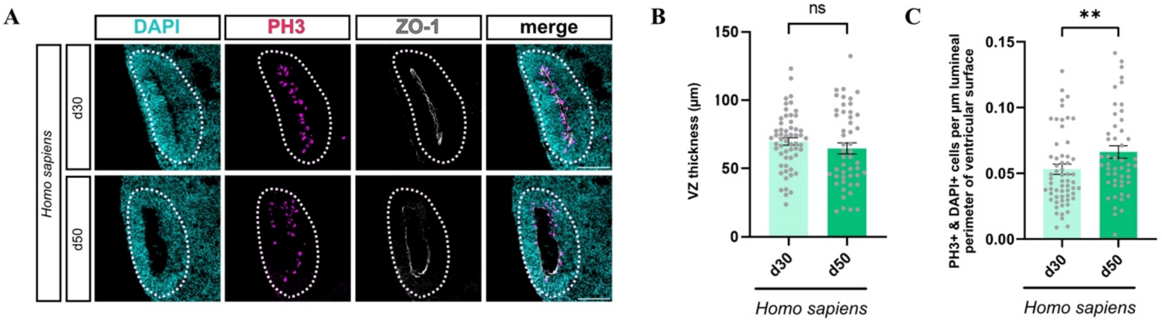


**Fig. S3: Human cerebral organoids maintain VZ thickness and consistent aRG mitotic activity during early neurogenesis.**

**(A)** Double immunofluorescence for PH3 (magenta) and ZO-1 (gray), combined with DAPI staining (cyan), in 30- (upper row) and 50 (lower row)-day-old human cerebral organoids. Dashed lines indicate VZ/basal region border. Scale bars, 100 µm. **(B)** Quantification of VZ thickness (µm) measured at four locations spaced 90° apart around a ventricle-like structure in 30- and 50-day-old human cerebral organoids. Data represent the mean of 57 ventricle-like structures from 16 human 30-day-old cerebral organoids, and 49 ventricle-like structures from 12 human 50-day-old cerebral organoids, generated from 3 independent human iPSC lines. Error bars, ±SEM; ns not significant: P=0,2714 (Mann-Whitney test). **(C)** Quantification of PH3+ mitotic cells at the ventricular surface, normalized to the luminal perimeter, in 30- and 50-day-old cerebral organoids. Data represents the mean of 60 ventricle-like structures from 16 human 30-day-old cerebral organoids, and 50 ventricle-like structures from 12 human 50-day-old cerebral organoids, generated from 3 independent human iPSC lines. Error bars, ±SEM; **P=0.0093 (Mann-Whitney test).

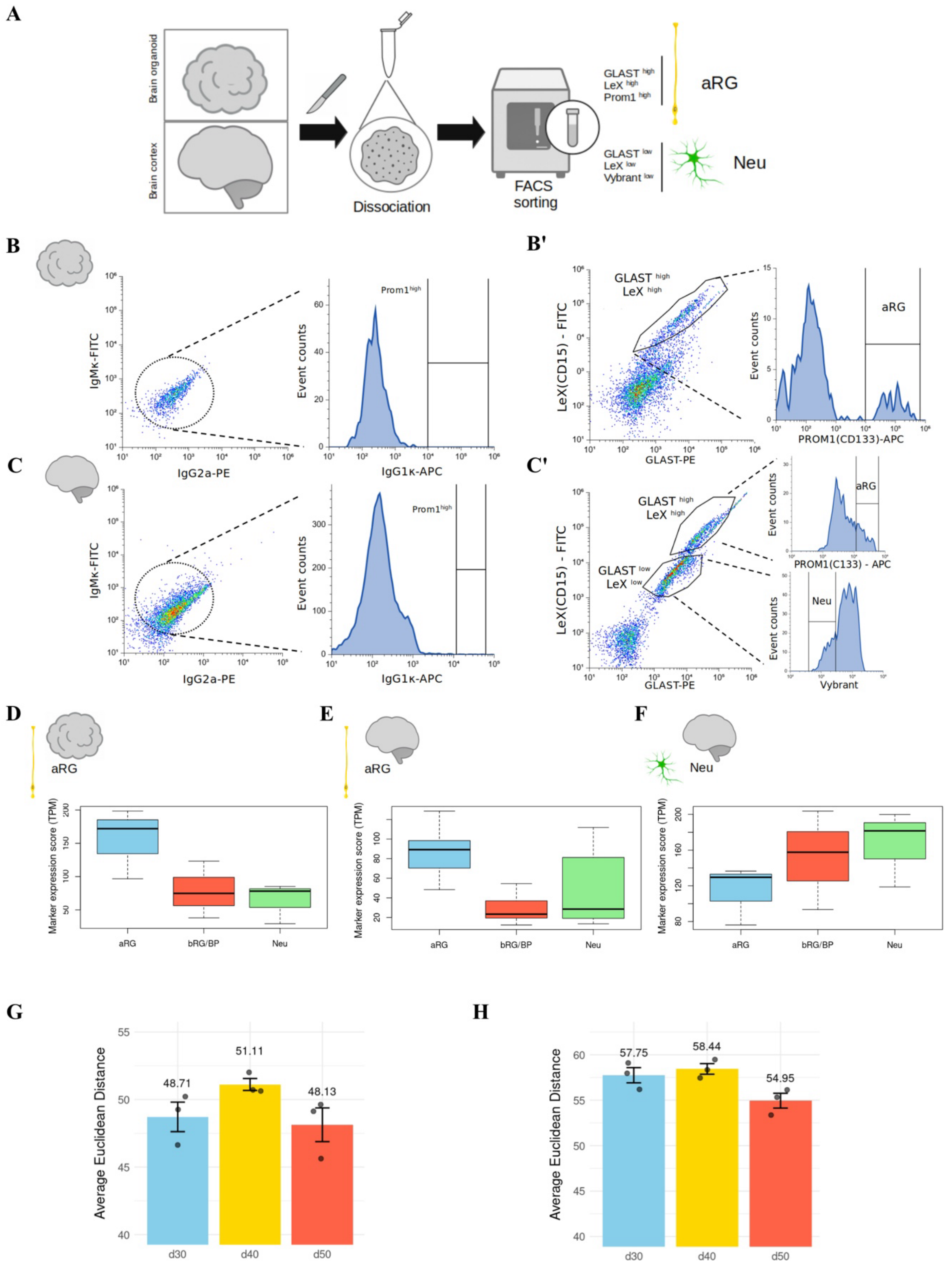
A
Brain organoid
Brain cortex
Dissociation
FACS sorting
GLAST high
LeX high
Prom1 high
aRG
GLAST low
LeX low
Vybrant low
Neu
B
B'
C
C'
D
E
F
G
H
Average Euclidean Distance
48.71
51.11
48.13
57.75
58.44
54.95
d30
d40
d50

**Fig. S4: FACS-enriched aRG from marmoset cerebral organoid show greater transcriptomic similarity to *in vivo telencephalic* aRG than to telencephalic neurons.**

**(A)** Schematic illustration of fluorescence-activated cell sorting (FACS) experimental design used to enrich aRG and neuronal populations from marmoset cerebral organoids (days 30, 40 and 50) and GD90 fetal marmoset telencephalic tissue. **(B, B')** FACS gating strategy to enrich aRG from day 30, 40 and 50 marmoset cerebral organoids using GLAST, LeX, and PROM1 antibodies. Gates were set using isotype control antibodies **(B)**. Apical radial glia were enriched based on high GLAST and LeX signals (left plot), and high PROM1 signal (right plot) **(B')**. **(C, C')** FACS gating strategy to enrich aRG and neurons from GD90 fetal marmoset telencephalic tissue using the same antibodies as in **(B, B')** and Vybrant DNA staining to quantify DNA content. Gates were set using isotype control antibodies **(C)**. Apical radial glia were enriched as described in **(B')**. Neurons were enriched based on low GLAST and LeX signals (left plot), and low Vybrant signal (right plot) corresponding to low DNA content indicative of the G1/G0 phase **(C′).** FACS sorting gates for GLAST, LeX, PROM1, and Vybrant staining in GD90 telencephalon to isolate aRGs and neurons. **(D-F**) Marker gene expression (in transcripts per million, TPM) of key aRG, bRG/bIP, and neuronal markers in FACS-sorted populations of marmoset cerebral organoid aRG. Expression is shown for three cell lines at days 30, 40, and 50 of organoid culture (n = 9). **(D)**, GD90 telencephalic tissue aRG (*n* = 3) **(E)**, and GD90 telencephalic tissue neurons (*n* = 3) **(F)**. **(G,H)** Average Euclidean distance between FACS-sorted aRG from 30- (blue), 40- (yellow) and 50 (red)-days-old marmoset cerebral organoids and aRG **(G)** or neurons **(H)** from GD90 fetal marmoset telencephalic tissue, based on expression data of 728 neurodevelopmental genes **(see Table S5)**. The distance was measured from each indicated stage average to every individual GD90 telencephalon (n=3 per time point).

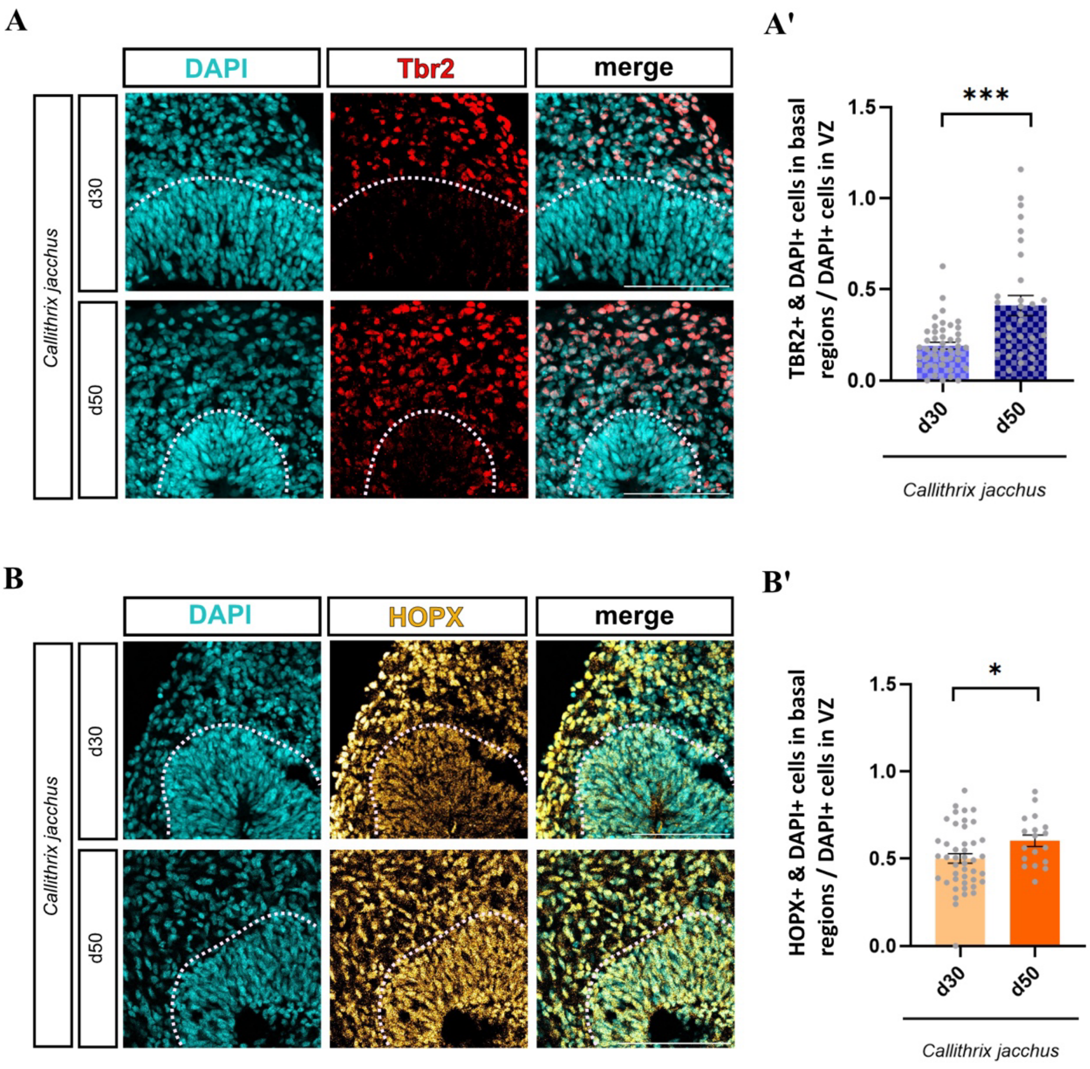


**Fig. S5: Marmoset AP delamination leads to increased abundance of BP cell types in cerebral organoids.**

**(A)** Immunofluorescence for Tbr2 (red), combined with DAPI staining (cyan) in 30- (upper row) and 50 (lower row)-day-old marmoset cerebral organoids. Dashed lines indicate VZ/basal region border. Scale bars, 100 µm. **(A')** Quantification of Tbr2+/DAPI+ double-positive cells in the basal region relative to DAPI+ VZ progenitors in the adjacent VZ region of 30- and 50-day-old marmoset cerebral organoids. Data represent the mean of 44 radial units from 19 marmoset 30-

day-old cerebral organoids, and 31 radial units from 23 marmoset 50-day-old cerebral organoids, generated from 3 independent marmoset iPSC lines. Error bars, ±SEM; ***P=0.0004 (Mann-Whitney test). **(B)** Immunofluorescence for HOPX (orange), combined with DAPI staining (cyan), in 30- (upper row) and 50 (lower row)-day-old marmoset cerebral organoids. Dashed lines indicate VZ/basal region border. Scale bars, 100 µm. **(B')** Quantification of HOPX+/DAPI+ double-positive cells in the basal region compared to DAPI+ VZ progenitors in the adjacent VZ region of 30- and 50-day-old marmoset cerebral organoids. Data represent the mean of 44 ventricle-like structures from 15 marmoset 30-day-old cerebral organoids, and 18 ventricle-like structures from 10 marmoset 50-day-old cerebral organoids, generated from 3 independent marmoset iPSC lines. Error bars, ±SEM; *P=0.0377 (unpaired t-test).

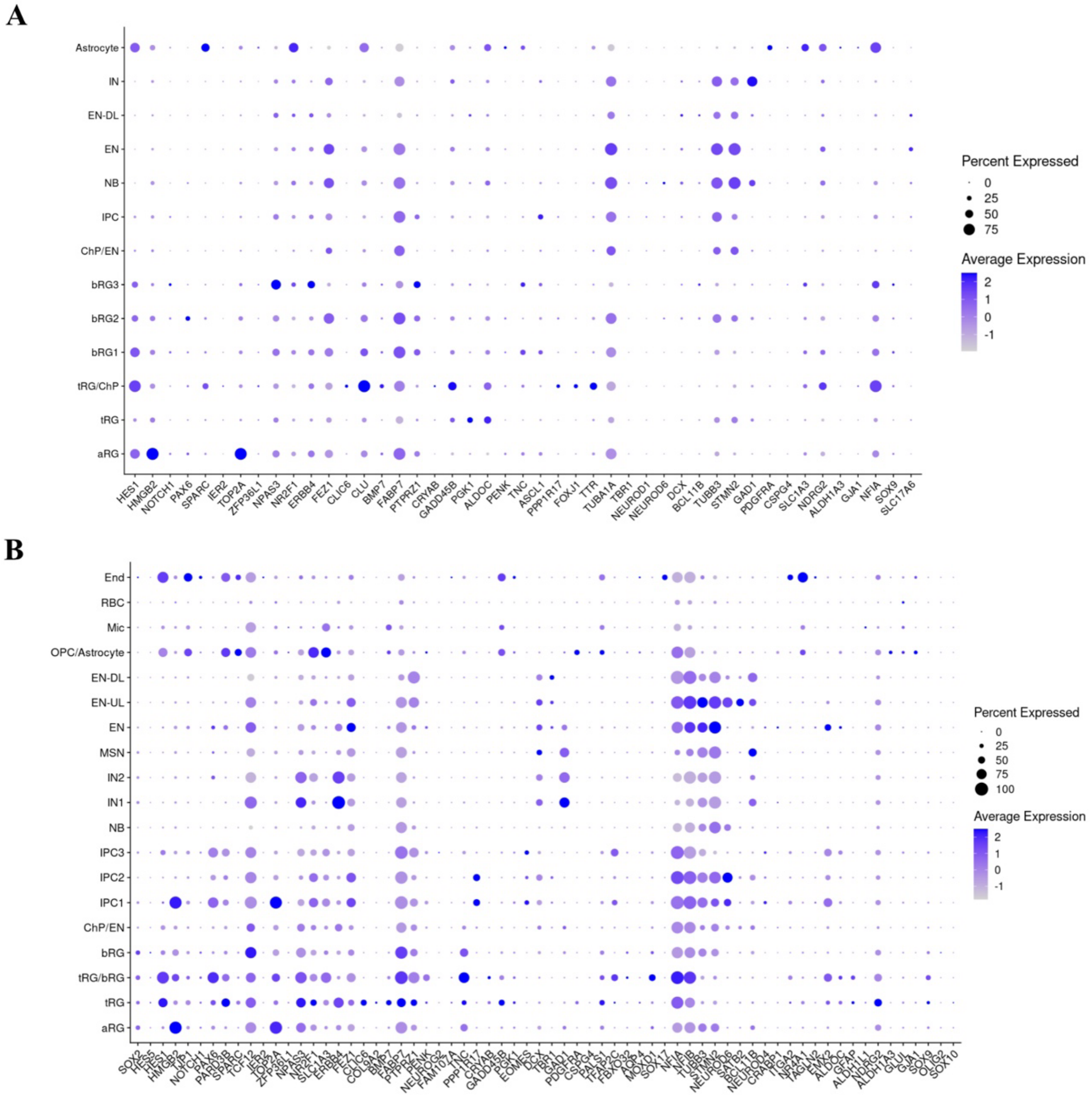


**Fig. S6: Marker gene expression confirms the identity of annotated cell types in scRNA-seq data from marmoset day 50 cerebral organoids and GD90 telencephalic tissue.**

**(A-B)** Dot plots of SCTransform-normalized scRNA-seq expression data showing average expression levels (color) and the percentage of cells expressing selected marker genes as indicated (dot size) in day 50 marmoset cerebral organoids **(A)** and GD90 fetal marmoset telencephalic tissue **(B)**.

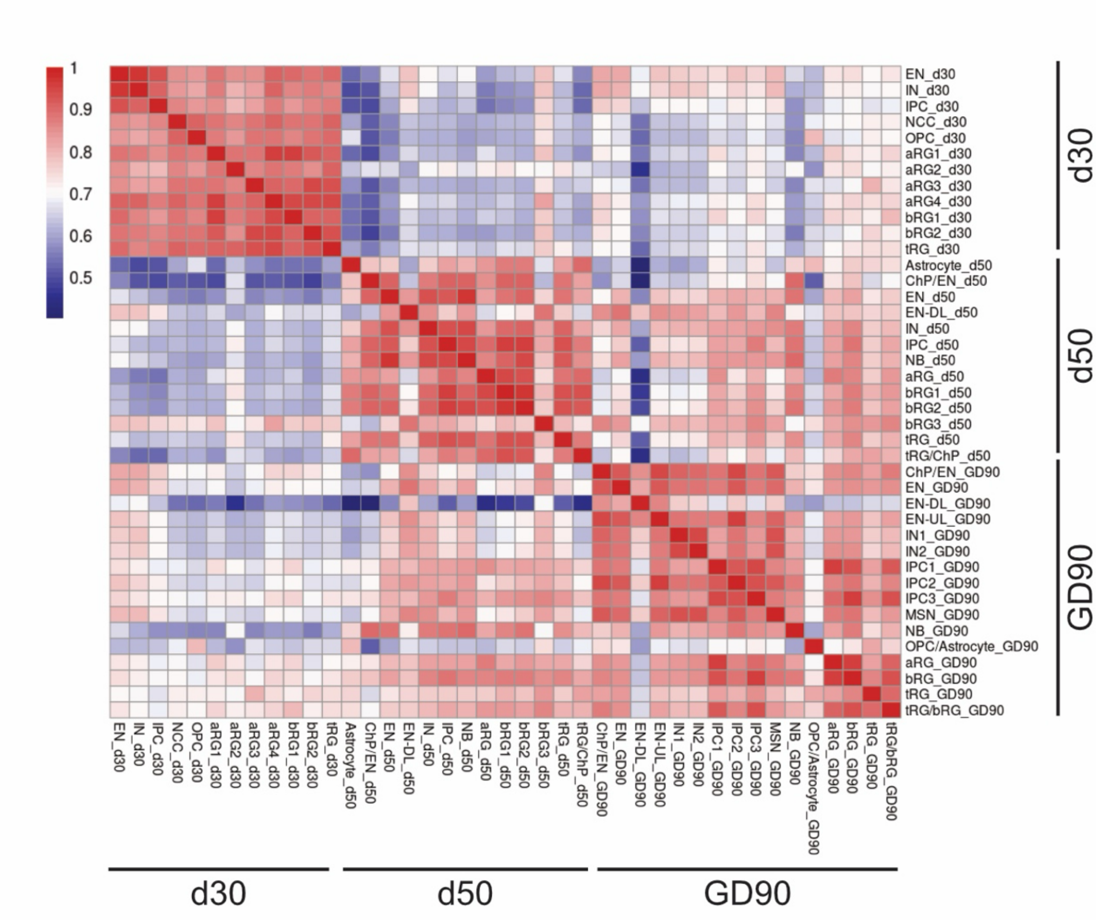


**Fig. S7: The highest cross-sample correlations are observed between day 50 marmoset cerebral organoid clusters and GD90 fetal marmoset telencephalic.**

Heatmap of Pearson correlation matrix of scRNA-seq clusters for all marmoset scRNA-seq samples (day 30 and day 50 cerebral organoids and GD90 telencephalic tissue).
